# Fucoidan-Copper Nanoparticles to Potentiate Synergistic Cancer Cell Cuproptosis and Immunotherapy

**DOI:** 10.1101/2024.10.20.619277

**Authors:** Hao-Hong Chen, Xiao-hui Pang, Qian-hui Wang, Ziye Chen, Li-li Zhuang, Jia-yuan Luo, Qian-xi Zheng, Rui-fang Zhong, Xiao-mei Zhan, Li Yang, Liang Zhu, Jian-Guo Jiang

## Abstract

Cuproptosis, a newly characterized form of regulated cell death initiated by copper binding to lipoylated components of the tricarboxylic acid cycle, presents a promising target for cancer therapy. Here, we report the development of fucoidan-copper nanoparticles (Fu-Cu) that exploit this mechanism to selectively induce cytotoxicity in HuH-7 liver cancer cells. The Fu-Cu was synthesized using fucoidan, a sulfated polysaccharide with inherent anticancer properties, as a natural nanocarrier for copper ions. Characterization confirmed successful copper incorporation and the formation of stable nanoparticles. Fu-Cu treatment enhanced intracellular copper levels and oxidative stress, triggering cuproptosis mediated by mitochondrial carrier homolog 2. Knockout of ferredoxin 1 in HuH-7 cells mitigated the cytotoxic effects, underscoring its critical role in copper-induced cell death. In *vivo* studies using a subcutaneous tumor model in BALB/c nude mice demonstrated that Fu-Cu effectively inhibited tumor growth and stimulated antitumor immunity, evidenced by increased infiltration of T cells, natural killer cells, and macrophages within the tumor microenvironment. These findings highlight Fu-Cu as a novel therapeutic strategy for HCC, leveraging the mechanism of cuproptosis and immune activation to suppress tumor progression.

## Main

Hepatocellular carcinoma (HCC) is the fifth most common malignant tumor worldwide and ranks as the third leading cause of cancer-related deaths, with a five-year survival rate of approximately 7%^1,2^. The progression of HCC involves a multistep process, transitioning from chronic inflammation and cirrhosis to primary and metastatic carcinoma^3^. The aggressive nature of the disease and frequent late diagnoses highlight the urgent need for novel and highly effective treatments.

Copper is essential for cellular functions but becomes toxic when its homeostasis is disrupted. Elevated copper levels have been consistently observed in the serum and tumor tissues of cancer patients compared to healthy individuals^4^. Dysregulation of copper homeostasis can lead to cellular toxicity, and recent studies have identified a unique form of regulated cell death termed “cuproptosis”^5–7^. This mechanism is distinct from apoptosis, necrosis, pyroptosis, and ferroptosis, and is initiated by the direct binding of copper ions to lipoylated components of the tricarboxylic acid (TCA) cycle within mitochondria^5,8–11^. The exploitation of cuproptosis presents a promising avenue for targeted cancer therapy, particularly in cancers with aberrant copper metabolism.

Copper has demonstrated therapeutic effects against cancer cells, playing a crucial role in various physiological processes, including enzymatic reactions and mitochondrial respiration. However, its therapeutic potential is limited by the narrow margin between its beneficial and toxic doses^4^. Fucoidan (Fu), a sulfated polysaccharide derived from brown seaweed, has garnered significant attention for its anticancer properties, including inhibition of tumor growth and metastasis, as well as modulation of the immune response^12–14^. Given the complementary properties of these two compounds, the development of fucoidan-based copper nanoparticles for cancer therapy presents a promising strategy. This approach could leverage cuproptosis to target cancer cells, while fucoidan’s drug delivery capability and tumor-specific recognition could minimize toxicity to normal cells, enhancing the therapeutic index.

Here, we developed fucoidan-copper nanoparticles (Fu-Cu) for HCC, focusing on their synthesis, characterization, and potential efficacy. Fu-Cu leverage fucoidan as a natural carrier to enhance copper ion delivery, inducing selective liver cancer cell death through cuproptosis via mitochondrial dysfunction and proteotoxic stress. Additionally, the immunomodulatory properties of fucoidan may boost antitumor immunity, offering a dual therapeutic approach. By exploring the interactions between copper ions and cellular components involved in cuproptosis, as well as the roles of key proteins such as mitochondrial carrier homolog 2 (MTCH2)^15–17^ and ferredoxin 1 (FDX1)^18–20^, we aim to elucidate the underlying mechanisms of Fu-Cu-induced cytotoxicity and consider the implications of these findings for the development of targeted therapies that can overcome the limitations of current treatments for HCC.

## Results

### Characterization of Fu-Cu

Fucoidan, a polysaccharide extracted from brown seaweed, was selected for its notable anticancer activity, high biocompatibility, and low toxicity to serve as a nanocarrier for copper. Fu-Cu nanoparticles were successfully synthesized by chemical synthesis method and using CuCl_2_ as the copper source (Figure 1a and Figure S1). The freeze-dried Fu-Cu powder appeared yellow-green to light blue, with a loose flocculent texture and no noticeable odor (Figure 1b). The identification reaction for Cu²⁺ ions showed no significant color change in the liquid outside the Fu-Cu dialysis bag, while the liquid inside the bag appeared light blue (Figure 2c). Upon the addition of strong acid, the solution turned a distinct deep blue. These results indicate that the majority of copper in the prepared Fu-Cu solution is chemically bonded to the polysaccharide, rather than being a simple mixture (Figure 1c).

**Figure 1.**
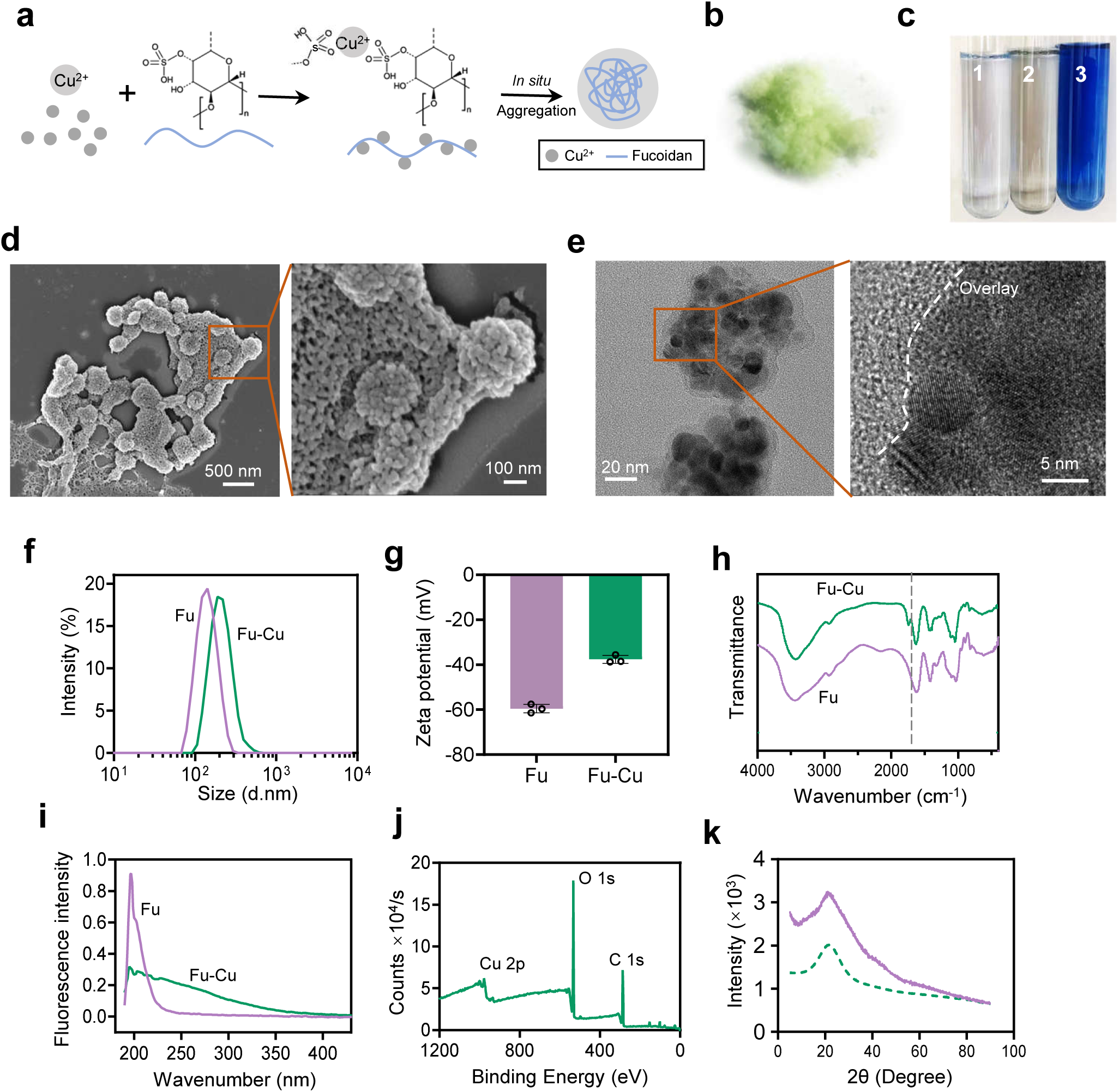
Synthesis and characterization of Fu-Cu nanoparticles. **a.** Schematic illustration of the in situ formation of Fu-Cu via coordination between Cu²⁺ and fucoidan, leading to aggregation. **b.** Digital image of freeze-dried Fu-Cu powder, showing a yellow-green to light blue, flocculent appearance. **c.** Copper ion identification assay: 1) liquid from outside the Fu-Cu dialysis bag (no color change), 2) liquid from inside the dialysis bag (light blue), 3) liquid from inside the bag with the addition of 1 M HCl (deep blue), indicating copper ion binding to fucoidan. SEM **(d)** and TEM **(e)** images of Fu-Cu showing spherical morphology with an average size of ∼200 nm. The higher magnification TEM image highlights the encapsulation of smaller particles within the larger structures. **f.** Particle size distribution of Fu and Fu-Cu measured by Nanoparticle Tracking Analysis, showing an increase in size upon copper incorporation. **g.** Zeta potential measurements of Fu and Fu-Cu, demonstrating a reduction in surface charge after copper binding. **h.** FTIR spectra of Fu and Fu-Cu, highlighting characteristic sulfated polysaccharide peaks and the emergence of a new peak at 1735 cm⁻¹, indicating O-acetyl group formation. **i.** UV-vis absorption spectra of Fu and Fu-Cu, with Fu-Cu showing a broader absorption range and distinct spectral shifts compared to Fu. **j.** XPS spectrum of Fu-Cu confirming the presence of Cu²⁺ along with C, O, and N elements. **k.** XRD patterns of Fu and Fu-Cu, showing the amorphous nature of Fu-Cu with a broad peak around 2θ = 21°. Data are presented as mean values ± SD, error bars indicate standard deviations (n =3, biologically independent samples).

**Figure 2.**
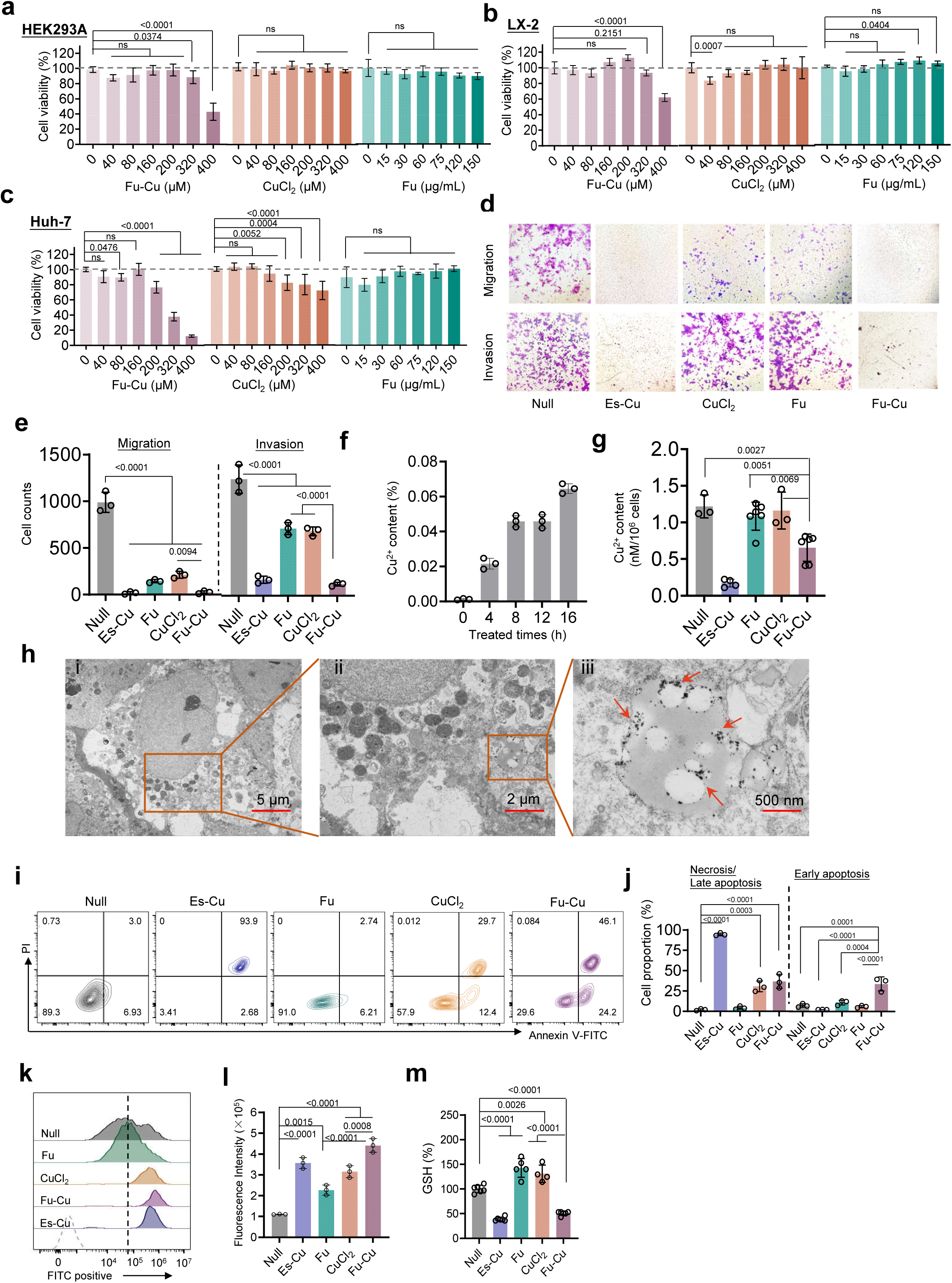
Cytotoxic effects and antitumor mechanisms of Fu-Cu. Cell viability assays of HEK-293A (**a**) and LX-2 (**b**) and HuH-7 cells (**c**) treated with Fu, CuCl₂, and Fu-Cu at various concentrations. Fu significantly promoted the proliferation of LX-2 cells, while CuCl₂ exhibited minimal cytotoxicity toward normal cells. Fu-Cu showed strong inhibition of cell viability in HuH-7 cells, demonstrating superior anticancer activity compared to Fu and CuCl₂. **d.** Representative Transwell assay images showing migration and invasion of HuH-7 cells under different treatments. **e.** Quantification of Transwell assay results showing that Fu-Cu significantly inhibited both migration and invasion compared to Fu or CuCl₂ alone (*P* < 0.0001). **f.** Total copper content in HuH-7 cells treated with Fu-Cu, measured by ICP-MS over different time points. **g.** Cu²⁺ content in HuH-7 cells measured using a cell copper assay kit. Fu-Cu significantly reduced Cu²⁺ levels while increasing total intracellular copper, indicating reduction of Cu²⁺ to Cu⁺. **h.** TEM images of HuH-7 cells treated with Fu-Cu. Red arrows indicate Fu-Cu internalized via endocytosis. Flow cytometry analysis (**i**) and quantification (**j**) of apoptotic and necrotic populations in HuH-7 cells after treatment, showing increased early apoptosis in Fu-Cu-treated cells compared to CuCl₂. Flow cytometry profiles (**k**) and fluorescence intensity quantification (**l**) of ROS in HuH-7 cells, showing significantly elevated ROS levels after Fu-Cu treatment, surpassing those induced by CuCl₂ and Es-Cu. **m.** Relative GSH content in HuH-7 cells under various treatments. Fu-Cu significantly depleted GSH levels, contributing to oxidative stress and apoptosis. Data are presented as mean values ± SD, error bars indicate standard deviations (n =3, biologically independent samples).

Scanning electron microscopy (SEM) and transmission electron microscopy (TEM) revealed that Fu-Cu has an approximately spherical structure with an average particle size around 200 nm (Figure 1d and 1e). Multiple small particles were present within Fu-Cu, which were subsequently encapsulated into larger spherical structures. This may be due to the sulfate groups on Fu binding with multiple Cu²⁺, forming fucoidan-copper nanoparticles. The SEM images also showed that multiple smaller particles were encapsulated within larger spherical structures, which could be attributed to the binding of multiple Cu^2+^ to fucoidan sulfate groups. The surface of Fu-Cu was relatively rough, with no notable aggregation observed, indicating enhanced stability attributed to the hydrophobic properties of fucoidan.

The average particle size of Fu was measured at 139.96 ± 2.65 nm with a zeta potential of -59.5 ± 1.95 mV. In contrast, Fu-Cu exhibited a larger average particle size of 257.23 ± 3.91 nm and a reduced zeta potential of -37.6 ± 1.73 mV. Both Fu and Fu-Cu displayed a unimodal size distribution with a PDI of less than 0.3, indicating a uniform particle size distribution (Figure 1f and 1g). The increase in particle size and the reduction in zeta potential upon copper incorporation are likely due to the binding of Cu^2+^ to the negatively charged sulfate groups in fucoidan, reducing the overall negative charge and leading to particle aggregation.

FTIR analysis of Fu and Fu-Cu both revealed a broad, strong absorption peak around 3438 cm^-1^, corresponding to hydroxyl group stretching^21,22^, and sharp absorption peaks near 1030 cm^-1^ and 820 cm^-1^, characteristic of sulfated polysaccharides^23^ (Figure 1h). Notably, a new strong absorption peak at 1735 cm^-1^ was observed in Fu-Cu, indicative of O-acetyl group formation^24^, likely resulting from the cleavage of glycosidic bonds and the formation of carboxyl groups upon copper binding. This structural modification supports the presence of a unique chemical structure in Fu-Cu. The UV-vis absorption spectra of Fu and Fu-Cu showed distinct differences, with both exhibiting strong absorption peaks around 200 nm (Figure 1i). However, Fu-Cu displayed a broader absorption range (200 ∼ 375 nm) and significantly lower absorbance at 200 nm compared to Fu. These differences indicate that Fu-Cu possesses a unique chemical structure, distinct from the original fucoidan, likely due to the binding of copper ions.

XPS analysis confirmed the presence of C, O, N, and Cu elements in Fu-Cu, with Cu 2p spectra showing spin-orbit splitting indicative of Cu(II) oxidation state^25^ (Figure 1j). XRD analysis revealed that Fu-Cu was amorphous, with a broad diffraction peak at 2θ = 21° and lower crystallinity compared to fucoidan (Figure 1k). The absence of distinct copper-related diffraction peaks further confirmed the amorphous nature of Fu-Cu. Inductively coupled plasma mass spectrometry (ICP-MS) analysis determined the copper content in the freeze-dried Fu-Cu powder to be 5.94% (Supplementary Table S1).

In summary, Fu-Cu was synthesized with copper ions binding to fucoidan, forming stable spherical structures. Characterization confirmed structural changes, copper incorporation, and an amorphous nature, with a copper content of 5.94%.

### Selective Anticancer Activity of Fu-Cu Nanoparticles in HuH-7 Cells

We assessed the cytotoxic effects of Fu, CuCl₂, and Fu-Cu on human normal cells (LX-2 hepatic stellate cells and HEK-293A kidney epithelial cells) and HuH-7 liver cancer cells to evaluate the differential toxicity and anticancer activity of Fu-Cu. The treatments exhibited concentration-dependent effects on cell viability across all cell lines tested (Figure 2a-c).

Fu significantly promoted the proliferation of LX-2 cells, with the highest viability observed at 133.12% for concentrations between 60 and 150 μg/mL (*P* < 0.0001). This suggests a potential proliferative effect of Fu on normal hepatic stellate cells. CuCl₂ did not significantly alter the viability of LX-2 or HEK-293A cells, indicating minimal cytotoxicity towards normal cells at the concentrations tested. In contrast, Fu-Cu exhibited a strong inhibitory effect on LX-2 cells at 400 μM, reducing viability to 71.25% (*P* < 0.0001). In HEK-293A cells, Fu-Cu inhibited cell proliferation at higher concentrations, with viabilities of 88.54% at 320 μM (*P* < 0.05) and 42.81% at 400 μM (*P* < 0.0001). These results indicate that while Fu-Cu has some cytotoxic effects on normal cells at elevated concentrations, it is relatively selective at lower concentrations. HuH-7 cells were significantly inhibited by CuCl₂ in a concentration-dependent manner, with the lowest viability at 60.09% at 400 μM (*P* < 0.0001). Fu-Cu further reduced HuH-7 cells viability, achieving below 20% viability at 400 μM, demonstrating superior anticancer activity compared to Fu and CuCl₂ individually (Figure 2c). This pronounced cytotoxicity towards cancer cells highlights the potential of Fu-Cu as an effective anticancer agent.

The anti-metastatic properties of Fu-Cu were evaluated by analyzing its impact on the migratory and invasive abilities of HuH-7 cells using wound healing and transwell invasion assays. Fu-Cu significantly inhibited both migration and invasion of HuH-7 cells (*P* < 0.0001), more effectively than Fu or CuCl₂ alone (Figure 2d-e). The extent of inhibition was comparable to that observed with Es-Cu (100 nM Elesclomol + 10 μM CuCl_2_), a known copper-based anticancer agent, indicating that Fu-Cu possess strong anti-metastatic properties in addition to its cytotoxic effects on cancer cells.

To elucidate the mechanism underlying the enhanced anticancer activity of Fu-Cu, we measured the intracellular levels of Cu²⁺ and total copper in HuH-7 cells using a cell Cu assay kit and inductively coupled plasma mass spectrometry (ICP-MS), respectively. As shown in Figure 2g, the Cu²⁺ content in the Fu-Cu group was significantly reduced compared to the Null group (*P* < 0.01), although it remained higher than in the Es-Cu group. The Fu and CuCl₂ groups did not exhibit significant differences from the Null group in Cu²⁺ content. Simultaneously, a time-course study using ICP-MS revealed that the total copper content in HuH-7 cells treated with Fu-Cu increased over time (Figure 2f). HuH-7 cells were incubated with Fu-Cu for 0, 4, 8, 12, and 16 hours, and the total copper content showed an upward trend with extended incubation periods.

These findings indicate that while intracellular Cu²⁺ content decreases, the total copper content increases, suggesting a significant rise in Cu⁺ content within HuH-7 cells. The decrease in Cu²⁺ alongside an increase in total copper implies that Cu²⁺ is being reduced to Cu⁺ intracellularly. TEM images further demonstrated that Fu-Cu enter the cytoplasm via endocytosis (Figure 2h). This internalization mechanism facilitates the accumulation of copper within the cells, contributing to the enhanced cytotoxic effects observed.

We characterized the mode of cell death induced by Fu-Cu in HuH-7 cells using Annexin V-FITC/PI staining followed by flow cytometry analysis. After 24 hours treatment, Fu-Cu significantly increased the proportion of late apoptotic/necrotic cells, with a notable rise in early apoptotic cells compared to CuCl₂ (Figure 2i-j and Figure S2). The early apoptotic cells population in the Fu-Cu group reached 29.6%, significantly higher than in the CuCl₂ group. As the treatment duration increased from 18 to 24 hours, the proportion of early apoptotic cells nearly doubled, indicating a time-dependent enhancement of Fu-Cu-induced apoptosis. These results suggest that Fu-Cu induces apoptosis as a primary mechanism of cytotoxicity in HuH-7 cells.

To investigate the role of oxidative stress in Fu-Cu-induced cytotoxicity, we measured reactive oxygen species (ROS) levels and reduced glutathione (GSH) content in HuH-7 cells. Fu-Cu treatment led to a significant increase in ROS levels, surpassing those observed with CuCl₂ and Es-Cu (Figure 2k-l). This suggests that Fu-Cu enhances ROS generation, potentially through a copper-based Fenton reaction, contributing to oxidative damage and cell death^5,26,27^. Additionally, Fu-Cu significantly reduced GSH content in HuH-7 cells (*P*< 0.0001), similar to the effects observed in the Es-Cu group (Figure 2m). The depletion of GSH indicates a disruption of cellular redox homeostasis^28^, further contributing to the induction of apoptosis.

In summary, Fu-Cu demonstrated selective cytotoxicity, with minimal effects on LX-2 and HEK-293A cells at lower concentrations, while significantly inhibiting the proliferation of HuH-7 cells, especially at higher concentrations. The enhanced anticancer activity of Fu-Cu is attributed to its ability to increase intracellular Cu⁺ levels, induce apoptosis, and promote oxidative stress specifically in cancer cells, highlighting its potential as a therapeutic agent for liver cancer.

### MTCH2 Functions in Cuproptosis

Recent research found that MTCH2 was a crucial mitochondrial outer membrane protein insertase, responsible for inserting tail-anchored proteins, signal-anchored proteins, and multi-pass transmembrane proteins into the mitochondrial outer membrane^15^. Dysfunction of MTCH2 was associated with various diseases, including impaired mitophagy, altered lipid homeostasis, and Alzheimer’s disease^15,16,29^.

Gene expression analysis comparing HCC patients with normal controls revealed a significant upregulation of several cuproptosis-related genes, including A*TOX1*, *CCS*, *COX17*, *SOD1*, *FDX1*, *MTCH2*, and *PDHA1*. Notably, *MTCH2* expression was markedly elevated in HCC tissues (*P* < 0.01, Figure 3a). Kaplan–Meier survival analysis indicated that patients with lower expression levels of *MTCH2* had significantly higher survival rates compared to those with elevated *MTCH2* expression (Figure 3b). Furthermore, although *MTCH2* expression remained consistently higher across different clinical stages of HCC compared to normal individuals, no significant differences were observed among the various stages (Figure 3c), suggesting that *MTCH2* upregulation is an early event in hepatocarcinogenesis. RT-qPCR analysis showed a significant upregulation of the *MTCH2* gene after Fu-Cu treatment, suggesting that the *MTCH2* gene is involved in Fu-Cu-induced cuproptosis (Figure 3d).

**Figure 3.**
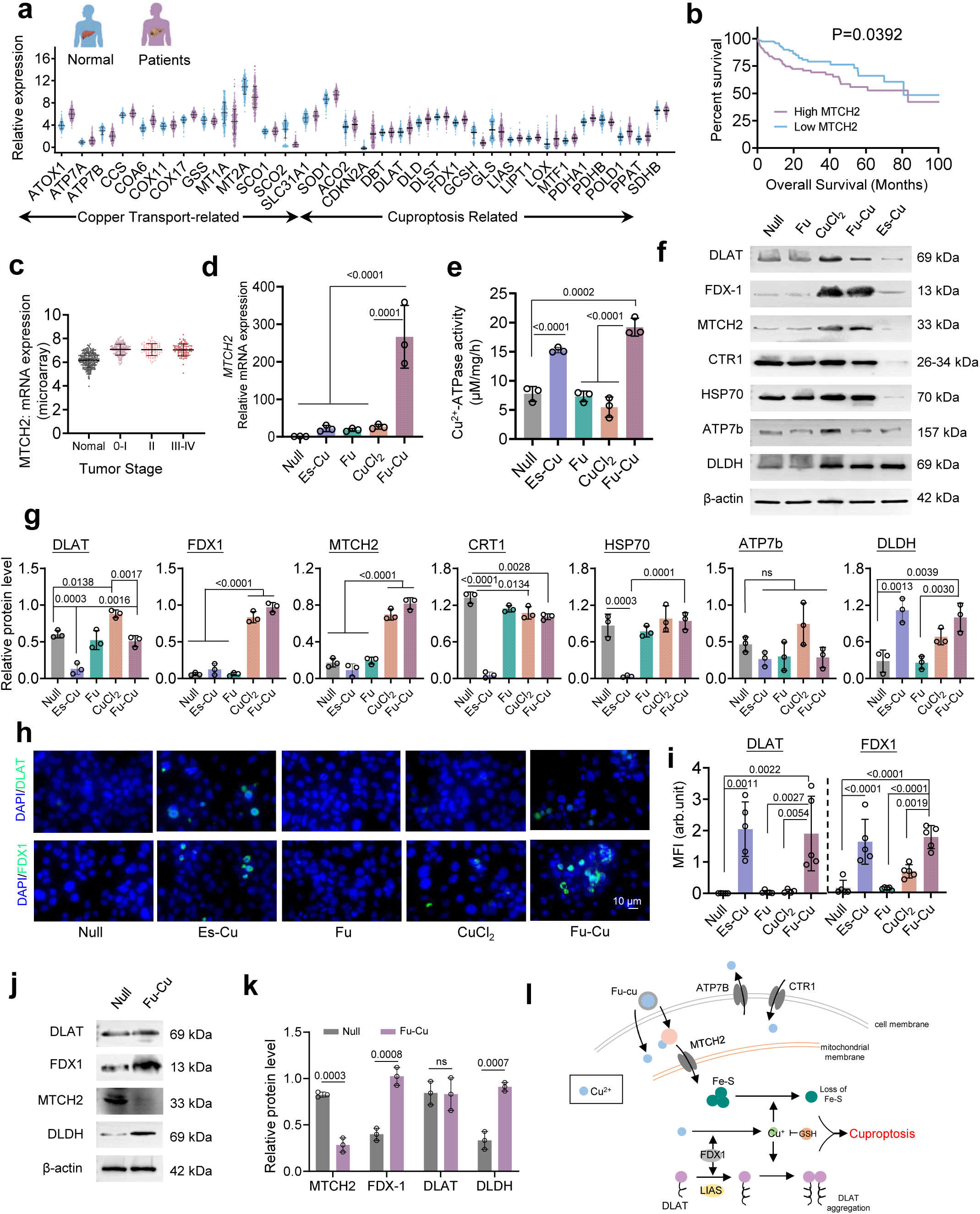
*MTCH2* involvement in copper-induced cuproptosis. **a.** Gene expression analysis comparing HCC patients with normal controls, showing upregulation of several copper transport- and cuproptosis-related genes, including *MTCH2*. **b.** Kaplan–Meier survival analysis showing that liver cancer patients with lower *MTCH2* expression have significantly higher survival rates than those with elevated *MTCH2* expression. **c.** *MTCH2* mRNA expression levels across different tumor stages, showing consistently higher expression in HCC patients compared to normal controls, without significant differences between stages. **d.** RT-qPCR analysis of *MTCH2* mRNA expression in HuH-7 cells treated with Es-Cu, Fu, CuCl₂, and Fu-Cu, showing significant upregulation of *MTCH2* in the Fu-Cu group. **e.** Cu²⁺-ATPase activity in HuH-7 cells, significantly elevated in the Fu-Cu and Es-Cu groups compared to the Null group, indicating enhanced copper metabolism. Western blot analysis (**f**) and quantification (**g**) of cuproptosis-related proteins (DLAT, FDX1, MTCH2, CRT1, HSP70, ATP7b, DLDH) in HuH-7 cells under different treatments. Fu-Cu treatment significantly upregulated FDX1, DLAT, DLDH, and MTCH2, while downregulating ATP7b. Immunofluorescence staining (**h**) and quantification (**i**) of FDX1 and DLAT in HuH-7 cells treated with Fu-Cu and Es-Cu, showing upregulation of these key cuproptosis markers. Scale bar, 10 μm. **j, k.** Western blot analysis (**j**) and quantification (**k**) of DLAT, FDX1, MTCH2, and DLDH in MTCH2-knockdown HuH-7 cells, showing reduced MTCH2 expression but compensatory upregulation of FDX1 and DLDH following Fu-Cu treatment. **l.** Schematic illustration of MTCH2’s role in facilitating the transport of Cu⁺-binding proteins into the mitochondria, promoting cuproptosis. Data are presented as mean values ± SD, error bars indicate standard deviations (n =3, biologically independent samples).

We assessed copper-ATPase activity in HuH-7 cells across different treatments to evaluate the impact of Fu-Cu on copper metabolism. The Fu-Cu and Es-Cu groups both exhibited a significant increase in copper-ATPase activity, reaching 19.67 ± 0.85 µM/mg/h and 15.42 ± 0.67 µM/mg/h respectively, compared to the Null group (8.23 ± 0.45 µM/mg/h, *P* < 0.001) (Figure 3e). In contrast, the Fu and CuCl₂ groups both showed no statistically significant difference in enzyme activity relative to the Null group (*P* > 0.05). These results suggest that Fu-Cu treatment enhances copper-ATPase activity, potentially influencing intracellular copper levels and promoting cuproptosis.

Western blotting analysis was performed to investigate the expression of key proteins involved in the cuproptosis pathway. Fu-Cu treatment markedly upregulated the expression of FDX1, DLAT, DLD, and MTCH2 proteins compared to the Null group (*P* < 0.01) (Figure 3f-g). Concurrently, ATP7B expression was significantly downregulated in the Fu-Cu group (*P* < 0.01), while the expression of CTR1 remained unchanged (*P* > 0.05). The downregulation of ATP7B, a crucial copper efflux transporter, indicates inhibited efflux of Cu⁺ ions, leading to intracellular copper accumulation. The unaltered CTR1 expression suggests that Cu⁺ uptake mechanisms are intact. These changes are associated with an intensified cuproptosis effect in HuH-7 cells under Fu-Cu treatment. In the CuCl₂ group, expression levels of FDX1, DLAT, DLD, and MTCH2 were also increased compared to the Null group (*P* < 0.05); however, ATP7B expression was significantly elevated (*P* < 0.01) (Figure 3f-g). The upregulation of ATP7B in this group facilitates the expulsion of excess Cu⁺ ions, thereby reducing intracellular copper toxicity and mitigating cuproptosis. In contrast, the Es-Cu group showed significant decreases in the expression levels of DLAT, FDX1, CTR1, HSP70, and MTCH2 relative to the Null group (*P* < 0.01) (Figure 3f-g). The pronounced reduction in CTR1 suggests impaired copper uptake, and the overall decrease in protein expression is likely due to extensive cell death induced by Es-Cu treatment after 24 hours, leading to diminished protein synthesis or increased degradation. The Fu group did not exhibit significant changes in protein expression levels compared to the Null group (*P* > 0.05), indicating that fucoidan alone does not substantially impact these pathways.

To further confirm the involvement of cuproptosis, we performed immunofluorescence staining for FDX1 and DLAT in HuH-7 cells. The Fu-Cu and Es-Cu treatment groups showed significant upregulation of FDX1 and DLAT proteins compared to the Null group (Figure 3h-i). The enhanced fluorescence intensity indicates increased expression of these key cuproptosis markers, supporting the occurrence of cuproptosis in cells treated with Fu-Cu and Es-Cu.

To elucidate the role of *MTCH2* in Fu-Cu-induced cuproptosis, we performed gene interference experiments to knock down *MTCH2* expression in HuH-7 cells (Figure S3). Following *MTCH2* knockdown, Fu-Cu treatment resulted in a significant inhibition of *MTCH2* expression (*P* < 0.01), as expected (Figure 3j-k). Interestingly, the expression of cuproptosis-related genes *FDX1* and *DLDH* were induced under these conditions (*P* < 0.05) (Figure 3j-k). These results suggest that while *MTCH2* expression is reduced, Fu-Cu can still promote the expression of other cuproptosis-related proteins, indicating a compensatory mechanism that partially sustains cuproptosis in the absence of *MTCH2*.

Our findings underscore the pivotal role of MTCH2 as a gatekeeper on the mitochondrial membrane in the process of Cu⁺-induced cuproptosis. MTCH2 facilitates the transport of Cu⁺-binding proteins into the mitochondria, a crucial step in the cuproptosis pathway (Figure 3l). The enhanced expression of MTCH2 in the Fu-Cu group correlates with increased Cu⁺ accumulation and heightened cuproptosis, highlighting its essential function in mediating copper-induced cell death in HuH-7 cells. The modulation of MTCH2 expression appears to significantly influence the extent of cuproptosis, emphasizing its potential as a therapeutic target for controlling copper-mediated cytotoxicity in liver cancer cells.

### *FDX1* Knockout Mitigates Fu-Cu-Induced Cuproptosis in HuH-7 Cells

FDX1 is a pivotal gene in the cuproptosis pathway, which plays a dual role in reducing Cu^2+^ to the more toxic Cu^+^ in the mitochondria and facilitating the binding of Cu^+^ to lipoylated proteins, leading to proteotoxic stress and cell death^5,8,30,31^. *FDX1* knockout in HuH-7 cells significantly reduced cellular respiration, leading to the accumulation of pyruvate and α-ketoglutarate and decreased protein lipoylation^5^. To explore the impact of *FDX1* deletion on copper toxicity, we generated HuH-7 *FDX1* knockout (KO) cells using CRISPR/Cas9 gene editing. This study aimed to determine whether the absence of *FDX1* affects the anticancer activity of Fu-Cu in HuH-7 cells.

The viability of HuH-7 *FDX1* KO cells was assessed following treatment with Fu, CuCl₂, and Fu-Cu (Figure 4a). Fu alone showed no significant inhibitory effect on HuH-7 *FDX1* KO cells, with higher concentrations promoting cell growth. In contrast, CuCl₂ exhibited a concentration-dependent response, enhancing cell viability at low concentrations and inhibiting it at higher concentrations. At 160 μM, the highest cell viability was observed (134.23%, *P* < 0.001), while at 400 μM, viability decreased to 84.50% (*P* < 0.05). Fu-Cu treatment resulted in a stronger inhibitory effect on HuH-7 *FDX1* KO cells than either Fu or CuCl_2_ alone, significantly reducing cell viability at 320 and 400 μM concentrations (48.20% and 24.55%, respectively, *P* < 0.0001). However, the inhibitory effect of Fu-Cu was less pronounced in HuH-7 *FDX1* KO cells compared to wild-type HuH-7 cells. Intracellular Cu^2+^ levels were significantly elevated in the CuCl_2_ and Fu-Cu groups compared to the Null group (Figure 4b, *P* < 0.0001), with Cu^2+^concentrations reaching 0.5037 nmol/10^6^ cells and 0.4135 nmol/10^6^ cells, respectively. In contrast, the Es-Cu group showed a marked decrease in Cu^2+^ levels (0.0284 nmol/10^6^ cells).

**Figure 4.**
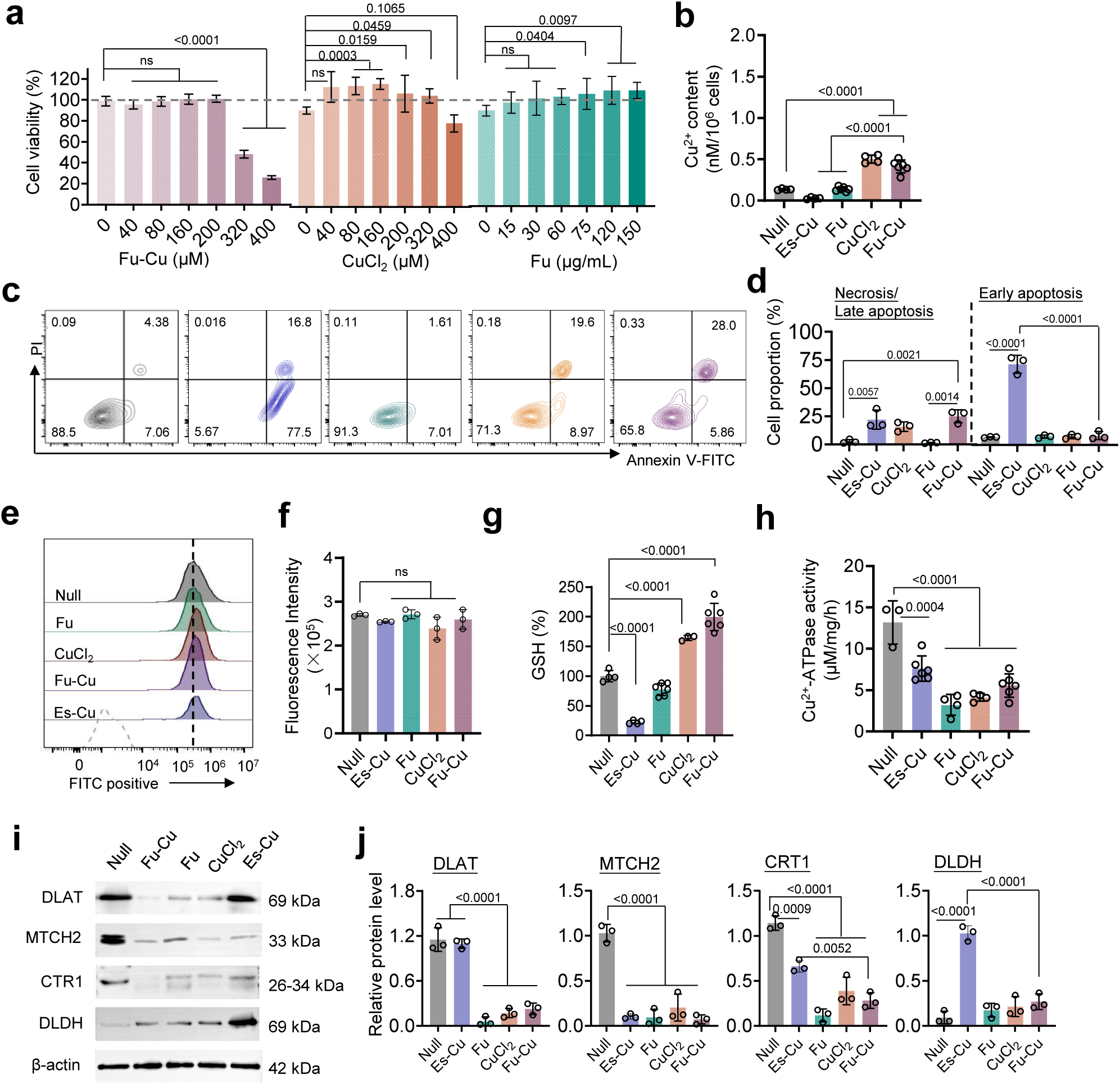
*FDX1* knockout mitigates Fu-Cu-induced cuproptosis in HuH-7 cells. **a.** Cell viability assays of HuH-7 *FDX1* knockout (KO) cells treated with Fu, CuCl₂, and Fu-Cu at different concentrations. *FDX1* KO cells exhibited reduced sensitivity to Fu-Cu compared to wild-type HuH-7 cells. **b.** Cu²⁺ content in *FDX1* KO HuH-7 cells measured using a cell copper assay kit, with elevated levels observed in the CuCl₂ and Fu-Cu groups. Flow cytometry analysis (**c**) and quantification (**d**) of necrosis/late apoptosis and early apoptosis in *FDX1* KO HuH-7 cells after treatment, indicating a reduction in Fu-Cu-induced apoptosis compared to wild-type cells. Flow cytometry profiles (**e**) and quantification (**f**) of ROS levels in *FDX1* KO HuH-7 cells under different treatments, showing no significant differences in ROS levels across treatments. **g.** Relative GSH content in *FDX1* KO HuH-7 cells, with elevated GSH levels in the Fu-Cu and CuCl₂ groups, suggesting GSH binding to copper may mitigate toxicity. **h.** Cu²⁺-ATPase activity in *FDX1* KO HuH-7 cells, with reduced activity across all treatment groups, particularly in the Fu group. Western blot analysis (**i**) and quantification (**j**) of cuproptosis-related proteins (DLAT, MTCH2, CTR1, DLDH) in *FDX1* KO HuH-7 cells, showing reduced expression of these proteins in Fu, CuCl₂, and Fu-Cu-treated cells. β-actin was used as a loading control. Data are presented as mean values ± SD, error bars indicate standard deviations (n =3, biologically independent samples).

Interestingly, despite these changes in copper levels, there were no significant differences in ROS levels between the treated groups and the Null group, with average fluorescence intensities around 250,000 (Figure 4e and 4f). This suggests that *FDX1* deletion reduces the extent of copper-induced mitochondrial dysfunction and cell death, thereby maintaining stable ROS levels. The Fu-Cu and CuCl_2_ treatments significantly increased GSH levels in HuH-7 *FDX1* KO cells (Figure 4g, *P* < 0.001), whereas the Es-Cu group exhibited a significant decrease in GSH levels (*P* < 0.001). In terms of copper-ATPase activity, all treatment groups, including Fu, CuCl_2_, and Fu-Cu, showed a significant reduction, with the Fu group displaying the lowest activity (3.21 µM/mg/h, Figure 4h). The increase in GSH levels in the Fu-Cu group suggests that GSH may bind copper, reducing intracellular copper content and thereby mitigating cuproptosis.

The deletion of *FDX1* significantly reduced the expression of MTCH2, CTR1, DLAT, and DLDH in the Fu, CuCl_2_, and Fu-Cu groups (Figure 4i and 4j). Notably, DLAT expression did not decrease in the Es-Cu group compared to the Null group. These findings indicate that Fu-Cu’s inhibitory effect on HuH-7 cells is closely associated with FDX1, as the deletion of FDX1 led to reduced DLAT and DLDH protein levels, correlating with a decrease in mitochondrial respiration. The absence of FDX1 also likely reduces protein lipoylation, preventing Cu^+^ binding to DLAT and thus alleviating proteotoxic stress and reducing Fu-Cu toxicity. Consequently, the HuH-7 *FDX1* KO cells exhibited higher viability under the same Fu-Cu treatment conditions compared to wild-type HuH-7 cells.

### Mechanistic Insights into Fu-Cu Toxicity in *C. elegans*

*Caenorhabditis elegans* possesses conserved copper transport and homeostasis mechanisms, which are pivotal for elucidating how dysregulation leads to cuproptosis^32^. In worms, MTCH1, the sole homolog of mammalian MTCH2, is suggested to regulate lipid homeostasis and promote apoptosis^33,34^. To further investigate the mechanisms underlying Fu-Cu-induced cuproptosis, we employed *C. elegans* as a model organism (Figure 5a-c).

**Figure 5.**
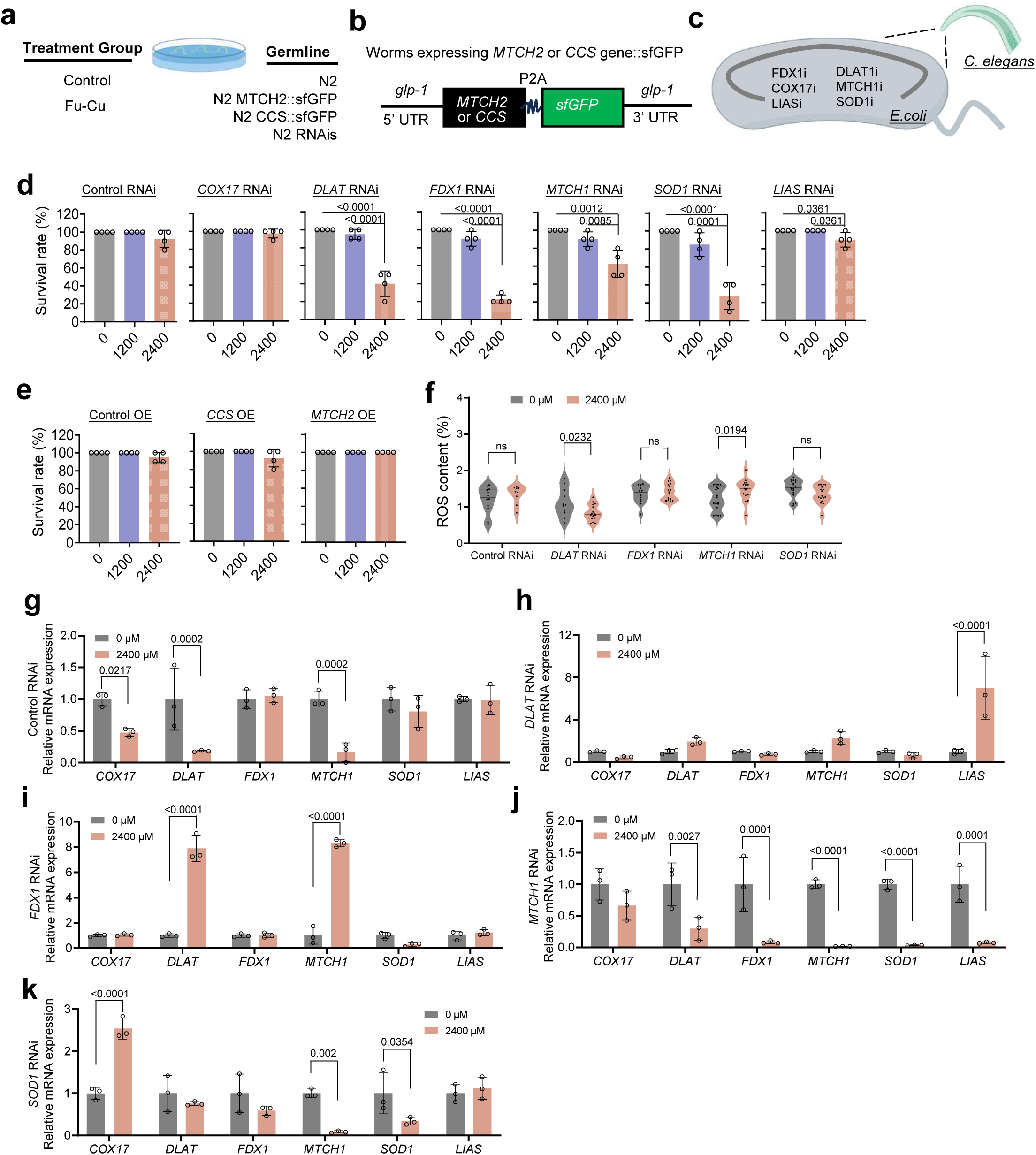
Investigation of Fu-Cu-induced cuproptosis in *C. elegans*. **a.** Schematic representation of the experimental setup, showing the treatment groups, including *C. elegans* strains treated with Fu-Cu and subjected to RNAi knockdown of cuproptosis-related genes. **b.** Diagram of transgenic *C. elegans* expressing *MTCH2* or *CCS* with GFP under the control of the *glp-1* promoter. **c.** Summary of key genes (*FDX1*, *COX17*, *DLAT*, *MTCH1*, *SOD1*, *LIAS*) targeted for RNAi in *C. elegans* and their role in cuproptosis. **d.** Survival rates of *C. elegans* RNAi mutants (*COX17*, *DLAT*, *FDX1*, *MTCH1*, *SOD1*, *LIAS*) under different concentrations of Fu-Cu (0, 1200, 2400 μM). *FDX1* and *SOD1* RNAi mutants exhibited significant sensitivity to Fu-Cu, with survival rates dropping to 22.5% and 27.5%, respectively, at 2400 μM Fu-Cu. **e.** Survival rates of *C. elegans* overexpressing human *MTCH2* or *CCS*, showing complete protection against Fu-Cu toxicity in the *MTCH2* overexpression group. **f.** ROS content in RNAi mutants (*DLAT*, *FDX1*, *MTCH1*, *SOD1*) after Fu-Cu treatment, with *DLAT* RNAi mutants showing a decrease in ROS levels and *MTCH1* RNAi mutants showing an increase. **g-l.** Gene expression analysis in RNAi mutants after Fu-Cu treatment. Fu-Cu exposure downregulated *COX17*, *DLAT*, and *MTCH2* in the Control RNAi group (**g**), while *LIAS* expression was upregulated in *DLAT* RNAi mutants (**h**). In *FDX1* RNAi mutants (**i**), Fu-Cu led to the upregulation of *DLAT* and *MTCH1*. Conversely, *MTCH1* RNAi mutants (**k**) showed broad downregulation of most genes, except *COX17*. *SOD1* RNAi mutants (**l**) showed upregulation of *COX17* and downregulation of *MTCH1* and *SOD1*. Data are presented as mean values ± SD, error bars indicate standard deviations (n =3, biologically independent samples).

Our study evaluated the toxicity of Fu-Cu in various gene knockdown mutants of *C. elegans*, focusing on *DLAT*, *FDX1*, *MTCH1*, and *SOD1*. And the human *MTCH2* and *CCS* genes were successfully introduced into *C. elegans* (Figure S4). Notably, RNAi-mediated knockdown of *FDX1* exhibited the highest sensitivity to Fu-Cu treatment, with survival rates plummeting to 22.5% under 2400 µM Fu-Cu exposure. Similarly, *SOD1* RNAi mutants showed significant susceptibility, with a survival rate of 27.5% at the same Fu-Cu concentration (Figure 5d and 5e). These findings underscore the potential role of ROS responses in copper-induced cell death, corroborated by the existing evidence linking ROS generation to copper cytotoxicity. Conversely, overexpression of human *MTCH2* provided complete protection against Fu-Cu toxicity, with survival rates reaching 100%.

To further elucidate the relationship between ROS and Fu-Cu toxicity, we assessed ROS levels in *DLAT*, *FDX1*, *SOD1*, and *MTCH1* RNAi mutants. Interestingly, *DLAT* RNAi mutants exhibited a decrease in ROS content post-Fu-Cu treatment compared to the untreated controls, whereas *MTCH1* RNAi mutants showed an increase in ROS levels under the same conditions (Figure 5f). These differential ROS responses suggest that specific genetic perturbations may either mitigate or exacerbate ROS-mediated copper toxicity.

Subsequent analysis of key gene expression in these RNAi mutants revealed diverse regulatory trends. In the Control RNAi group, Fu-Cu treatment led to the downregulation of *COX17*, *DLAT*, and *MTCH2*. In contrast, *LIAS* gene expression was significantly upregulated in *DLAT* RNAi mutants following Fu-Cu exposure. The *FDX1* RNAi group exhibited upregulation of *DLAT* and *MTCH1* genes (*P* < 0.0001), while *MTCH1* RNAi mutants displayed broad downregulation across most genes, except *COX17* (*P* ≤ 0.0027). In *SOD1* RNAi mutants, Fu-Cu treatment resulted in the upregulation of *COX17* (*P* < 0.0001) and downregulation of both *MTCH1* and *SOD1* genes (Figure 5g-h). These expression patterns highlight the complex genetic interactions underlying copper-induced cell death.

Contrary to expectations, active downregulation of key copper death-promoting proteins did not enhance resistance to copper toxicity; instead, it reduced survival rates among the *C. elegans* mutants. This suggests that copper-induced cell death may be a passive, rather than actively regulated, process. Furthermore, the downregulation of *MTCH2* was associated with the suppression of copper death-related gene expression, indicating a potential regulatory role for *MTCH2* in copper toxicity resistance mechanisms.

### Fu-Cu Trigger Antitumor Immunity and Cuproptosis in *vivo*

The immune system plays a crucial role in the surveillance and elimination of cancer cells, with immune cells such as T lymphocytes, natural killer (NK) cells, B cells, and macrophages contributing to antitumor immunity by recognizing and destroying malignant cells^35–37^. Understanding how anticancer therapies shape immune cell populations and functions is key to developing treatments that enhance immune responses. Fu-Cu showed anticancer effects in *vitro*, while in *vivo* studies are still needed to validate their impact on modulating the tumor microenvironment and enhancing immune-mediated tumor suppression.

Here, we utilized a BALB/c nude mouse model bearing subcutaneous HuH-7 liver cancer xenografts to evaluate the antitumor efficacy of Fu-Cu (Figure 6a). Dosages for Fu-Cu and other treatments were determined based on prior literature, preliminary experiments, and acute toxicity assessments in animals (Supplementary note 1 and Figure S5). Tumor volume measurements over the treatment period revealed that Fu-Cu significantly inhibited tumor growth compared to the PBS control group (*P* < 0.001) (Figure 6b). After 24 days of treatment, the tumors in the Fu-Cu group were markedly smaller than those in the PBS group (*P* = 0.0252), demonstrating the potent antitumor effect of Fu-Cu in *vivo* (Figure 6b-d).

**Figure 6.**
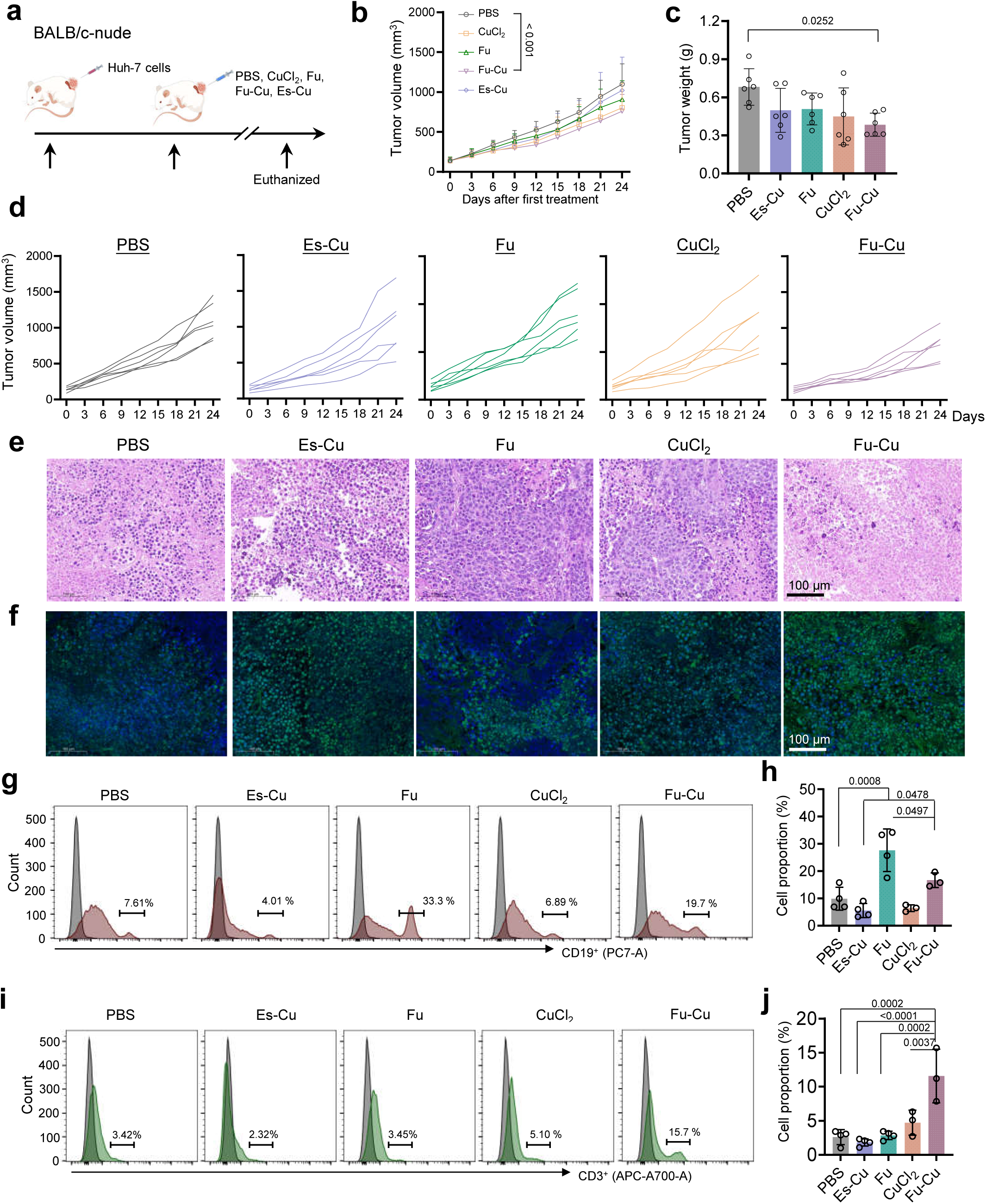
Fu-Cu enhances antitumor immunity and induces cuproptosis in *vivo*. **a.** Schematic illustration of the experimental setup using BALB/c nude mice bearing subcutaneous HuH-7 liver cancer xenografts. Mice were treated with PBS, CuCl₂, Fu, Fu-Cu, or Es-Cu over 24 days, with tumor volume and immune response analyzed. Arrows indicate intravenous injection time points. **b.** Tumor growth curves in mice over the 24-day treatment period. Fu-Cu significantly inhibited tumor growth compared to the PBS control group (*P* < 0.001). **c.** Final tumor weights measured at the endpoint, showing a significant reduction in tumor size in the Fu-Cu group (*P* = 0.0252). **d.** Individual tumor growth curves of mice across the different treatment groups, demonstrating the potent antitumor effect of Fu-Cu. **e.** Representative H&E staining images of tumor sections, showing cellular changes and increased cell death in Fu-Cu-treated tumors compared to controls. Scale bar, 100 μm. **f.** TUNEL staining images showing extensive apoptosis in tumors from Fu-Cu-treated mice. Scale bar, 100 μm. Representative flow cytometry profiles (**g**) and quantification (**h**) of CD19⁺ B cells in plasma, indicating elevated levels of CD19⁺ B cells in the Fu (33%) and Fu-Cu (19.7%) groups compared to other treatments. Representative flow cytometry profiles (**i**) and quantification (**j**) of CD3⁺ T cells in plasma, showing a significant increase in T cell populations in the Fu-Cu group (15.7%) compared to controls. Data are presented as mean values ± SD, error bars indicate standard deviations (n =3, biologically independent samples).

Histological analyses were performed to assess cellular changes within the tumor tissues. Hematoxylin and eosin (H&E) staining and TUNEL assays showed that the Fu-Cu-treated tumors exhibited significantly larger areas of cell death and apoptosis compared to the PBS group (Figure 6e 6f). These findings indicate that Fu-Cu effectively induces tumor cell death, contributing to the reduction in tumor size observed.

To investigate the impact of Fu-Cu on the immune system, we analyzed immune cell populations in the plasma and tumor tissues of treated mice. Flow cytometry of plasma samples revealed that the percentages of CD19^+^ B cells were higher in the Fu (33%) and Fu-Cu (19.7%) groups compared to other treatment groups (Figure 6g and 6h). Additionally, the Fu-Cu group showed a significant increase in CD3^+^ T cells, reaching 15.7%, which was substantially higher than in the other groups.

Immunofluorescence analysis of tumor tissue sections further demonstrated that the Fu and Fu-Cu groups had elevated levels of CD3^+^ T cells and CD56^+^ NK cells compared to other groups (Figure 7a and 7b). Moreover, the Fu-Cu group exhibited higher expression of F4/80^+^ macrophages within the tumor microenvironment (Figure 7c and 7d). These findings imply that Fu-Cu not only directly inhibits tumor growth but also stimulates the host immune system to mount a stronger antitumor response, contributing to its overall anticancer efficacy.

**Figure 7.**
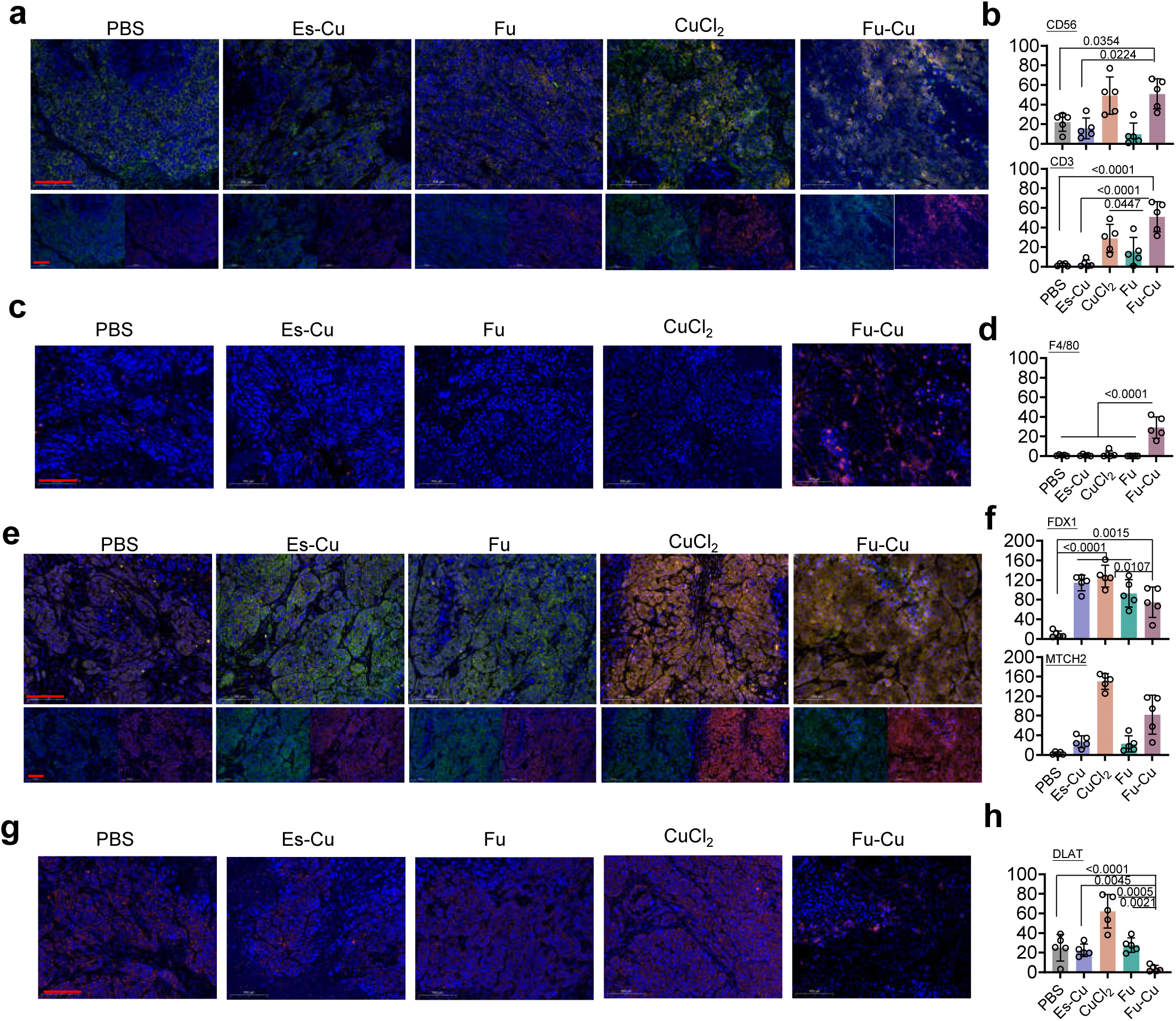
Fu-Cu enhances antitumor immunity and induces cuproptosis in *vivo*. Representative images (**a**) and quantifications (**b**) of immunofluorescence staining for CD56 (green), CD3 (red), and DAPI (blue) in tumor tissues from mice treated with PBS, Es-Cu, Fu, CuCl₂, and Fu-Cu. Fu-Cu-treated tumors showed significantly increased staining for CD56 and CD3, suggesting enhanced immune cell infiltration. Representative images (**c**) and quantifications (**d**) of immunofluorescence staining for F4/80 (red) and DAPI (blue), showing increased macrophage infiltration (F4/80) in Fu-Cu-treated tumors compared to other groups. Representative images (**e**) and quantifications (**f**) of immunofluorescence staining for FDX1 (green), MTCH2 (red), and DAPI (blue), highlighting elevated FDX1 and MTCH2 expression in Fu-Cu-treated tumors. Representative images (**g**) and quantifications (**h**) of immunofluorescence staining for DLAT (red) and DAPI (blue), showing significantly reduced DLAT expression in Fu-Cu-treated tumors, indicating the involvement of DLAT in Fu-Cu-induced cuproptosis. Scale bars= 100 μm. Data are presented as mean values ± SD, error bars indicate standard deviations (n =5, biologically independent samples).

We further explored the mechanism underlying the antitumor effects of Fu-Cu by examining the expression of proteins involved in the cuproptosis pathway within tumor tissues. Immunofluorescence staining was conducted for key cuproptosis-related proteins: FDX1, MTCH2, DLAT. The results revealed that MTCH2 expression was markedly upregulated in the CuCl₂ and Fu-Cu groups compared to other treatments (Figure 7e and 7f). The increased expression of MTCH2 suggests its significant involvement in cuproptosis within the tumor tissue.

The consistent upregulation of MTCH2 in both cellular and animal models underscores its importance in mediating the antitumor effects of Fu-Cu through the induction of cuproptosis. The activation of the cuproptosis pathway leads to the accumulation of toxic copper ions within cancer cells, promoting cell death and inhibiting tumor growth.

## Discussion

Our study demonstrates the promising anticancer effects of Fu-Cu, particularly in enhancing cytotoxicity through mechanisms involving cuproptosis and stimulation of the immune system. Despite these significant findings, several limitations must be addressed, and further research is required to fully realize the therapeutic potential of Fu-Cu.

While we identified MTCH2 as a critical mediator in this pathway, the specific interactions between copper ions and cuproptosis-related proteins remain unclear. Elucidating these molecular mechanisms could provide insights into how to enhance the efficacy of Fu-Cu and identify potential biomarkers for responsiveness to treatment. Moreover, the pharmacokinetics and biodistribution of Fu-Cu in *vivo* are not fully understood. Investigating the stability, delivery efficiency, and accumulation of Fu-Cu in tumor tissues versus normal tissues will be crucial for optimizing its therapeutic window and minimizing off-target effects.

Looking forward, our findings open several avenues for future research. One promising direction is the exploration of Fu-Cu in combination with immunotherapies. Given that Fu-Cu appears to enhance antitumor immune responses by increasing populations of T cells, NK cells, and macrophages, combining Fu-Cu with immune checkpoint inhibitors or adoptive cell therapies may synergistically improve treatment outcomes. Additionally, investigating the potential of Fu-Cu in other cancer types, particularly those with known copper metabolism dysregulation, could broaden its applicability. Identifying specific cancer subtypes that are more susceptible to cuproptosis could help tailor personalized treatment strategies.

Finally, to advance Fu-Cu towards clinical application, comprehensive preclinical studies focusing on scaling up production, evaluating safety profiles, and optimizing dosing regimens are essential. Clinical trials designed to assess the efficacy of Fu-Cu in HCC patients will be a critical step in translating our findings into a viable therapeutic option.

In conclusion, Fu-Cu represent a novel and promising therapeutic strategy for targeting copper metabolism in HCC. By inducing cuproptosis and stimulating the immune system, Fu-Cu has the potential to effectively inhibit tumor growth. Addressing the limitations of our study and pursuing the proposed future directions will be crucial in advancing Fu-Cu towards clinical application and expanding its utility across various cancer types.

## Methods

### Preparation of Fu-Cu

100 mg of fucoidan polysaccharide powder (Sigma) was accurately weighed, and after 80 mL of distilled water, it was placed in a magnetic stirrer. Stirred at 70℃ until fully dissolved, the volume was made up to 100 mL with distilled water, resulting in a 1 mg/mL solution of fucoidan. A 20 mM CuCl_2_ solution was prepared and filtered through a 0.45 μm membrane for later use. Under the action of a magnetic stirrer, 12.5 mL of CuCl_2_ solution (20 mM) was added to 100 mL of fucoidan solution. It was mixed thoroughly in the dark for 20 min. Then 12.5 mL of L-Ascorbic acid (80 mM) was added. After reacting at 50℃ for 4 hours, it was dialyzed with a 3500 kDa dialysis bag for 72 hours, and Fu-Cu was prepared. Reagents and chemicals used in this study are listed in Supplementary Table S2.

### Fu-Cu Characterization

Zeta Potential and Nanoparticle Size Analysis: The samples of Fu-Cu and Fu were diluted to an appropriate concentration, and the particles were ensured to be uniformly dispersed. The samples were then placed into Malvern ZETA potential sample dishes and Malvern plastic cuvettes. Subsequently, the Zeta potential distribution and particle size distribution were analyzed using a Nanoparticle Tracking Analysis (HORIBA).

Transmission Electron Microscopy and Scanning Electron Microscopy Analysis: The prepared samples of Fu-Cu were microscopically observed using Transmission Electron Microscopy (TEM, JEM-1400), focusing on the size, morphology, and uniformity of the samples. Specifically, the samples were dissolved in pure water, a few drops of anhydrous ethanol were added, and the mixture was sonicated for 5 minutes. The solution was then uniformly dropped onto a copper grid, dried under an infrared lamp for 2 minutes, and observed under an acceleration voltage of 80 kV. The morphology of Fu-Cu was also observed using Scanning Electron Microscopy (SEM, Hitachi S-800). A droplet of the sample suspension was placed onto a clean cover slip, which was then affixed to the sample stage using conductive double-sided tape. After gold sputtering, the sample was observed under the SEM at various magnifications ranging from 200 to 50,000 times.

Ultraviolet and Infrared Spectral Analysis: Sample solutions of appropriate concentrations were prepared and ultraviolet wavelength scans were conducted in the range of 190-840 nm, with pure water serving as a blank control. A quantity of 5 mg of the solid samples were meticulously mixed with potassium bromide (KBr) powder. After thorough grinding, the mixture was compressed using a pellet press. Subsequently, the infrared spectrum of the sample was recorded using a Fourier-transform infrared spectrometer (FTIR, Niolet iN10).

X-ray Analysis: A 50 mg dry powder sample of Fu-Cu was uniformly adhered to the sample stage using conductive double-sided tape, with dimensions less than 1×1 cm^2^ and a height less than 1.5 mm. The valence state of copper in the samples was analyzed using an X-ray Photoelectron Spectrometer (XPS, Thermo), and the crystal form of copper in the samples was analyzed using an X-ray single-crystal diffractometer (XRD, PANalytical B.V.). The diffraction angle range for the analysis was set between 10° and 90°.

Copper content in Fu-Cu: The copper content in the Fu-Cu samples were quantified using Inductively Coupled Plasma Mass Spectrometry (ICP-MS), with sample digestion facilitated by a graphite digester. Each sample was weighed and placed into graphite digester tubes. Subsequently, 2 mL each of nitric acid and hydrochloric acid were added to the sample, which were then allowed to stand at room temperature for approximately 15 minutes. The heating program for the graphite digester, specifically for copper-containing samples, was set as follows: 80°C for 10 minutes, 150°C for 10 minutes, and 200°C for 10 minutes. Upon the clarification of the digestion solutions, the samples were deemed fully digested. These solutions were then allowed to cool to room temperature. The digested samples were filtered into 100 mL volumetric flasks, diluted to the mark, and thoroughly mixed. The copper content was subsequently calculated using the given formula:

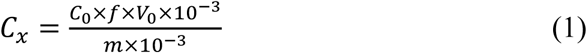

*C*_*x*_ = Final copper concentration (mg/kg); *C*_0_ = Detected copper concentration (mg/L); *f* = Dilution factor; *V*_0_ = Final digest volume (10 mL); m = Initial sample mass (g)

Cuprizone Colorimetric Method: Following 72 hours of dialysis, samples were collected from both inside and outside the Fu-Cu dialysis bags for copper ion identification. The dialysis water was replaced four times during this period, and the system was allowed to sit for 24 hours. Copper ion detection was performed using the cuprizone-based biscyclohexanone oxalyldihydrazone (BCO) color reagent, which forms a deep blue copper-ketone complex with Cu(II) ions in an alkaline environment. To prepare the BCO reagent, 0.5 g of cuprizone was dissolved in a mixture of 60 mL distilled water and 140 mL ethanol, with ultrasonic assistance to promote dissolution. For the assay, three glass test tubes were prepared. Tube 1 contained 10 mL of the liquid from outside the Fu-Cu dialysis bags, tube 2 contained 10 mL of the liquid from inside the dialysis bags, and tube 3 contained 5 mL of the liquid from inside the dialysis bags. To tube 3, 5 mL of 1 M HCl was added. Subsequently, 1 mL of ammonia and 1 mL of BCO color reagent were added to each tube. The tubes were incubated at room temperature for 15 minutes, and color changes were observed as an indication of copper ion presence.

### Cell Line Culture

The HuH-7, LX-2, and HEK293A cell lines were cultured in Dulbecco’s Modified Eagle Medium (DMEM) as a basal medium for cell growth and pseudoparticle production, supplemented with 10% (v/v) Gibco Fetal Bovine Serum (FBS) and 1% (v/v) Gibco Penicillin-Streptomycin (10,000 U/mL) when indicated. They were incubated at 37°C in a humidified 5% CO_2_ atmosphere. The HuH-7 and LX-2 cell lines were purchased from Procell (Wuhan, China).

### Genomic Manipulation of HuH-7 cell line

We employed a gene knockout approach to systematically in the HuH-7 cell line. Detailed information on the gRNAs and landing pads can be found in the primer list provided in Supplementary Table S3. A two-plasmid system was employed for genomic editing, consisting of a gRNA expression plasmid (pCKK001 SpCas9_gRNA, Zeocin) and the CRISPR/Cas9 expression plasmids (pCKK002 SpCas9, Neo), from Jiang Lab collection (Supplementary Table S4). The individual gRNAs were expressed using CMV promoter and bGH Terminator. A Cas9 expression cassette was constructed using the CMV promoter and hGH Terminator. For plasmid transformation, the CRISPR/Cas9 and gRNA expression plasmids were digested using *BsmB*I or *Bbs*I enzymes, respectively.

The design of all gRNAs was carried out using Benchling’s CRISPR Design Tool. Supplementary Table S3 provides a list of all the gRNAs used in this study. It was ensured that the 20 bp target sequences did not contain internal *Bsa*I, *BsmB*I, or NotI restriction sites to facilitate downstream cloning and transformation. The knockout gRNA assemblies were set up in 10 µL volume reactions, as follows: 1 μL primer (100 μM), 1 μL 10 × T4 DNA ligase buffer (Promega), 0.5 μL T4 PNK (NEB), and 7.5 μL H_2_O. After mixing, the mixture was incubated for 1 hour at 37°C. The sense and antisense oligonucleotide reactions (each 10 μL) were combined and brought to 200 μL with H_2_O. The oligonucleotides were annealed using slowly decreasing temperature cycles: 96°C for 6 minutes, followed by a ramp down of 0.1°C/s to 20°C, and a 20°C hold. A standard Golden Gate assembly reaction was performed, using 10 μL of the resulting reaction for ligation into the pCKK001 vector. The donor DNA was generated by cloning the sequence into the pDTK001 part entry vector^38,39^, which contains 500 bp of homology flanking the insert for multiplexed editing.

### Cell Knockout Transfection and Monoclonal Screening

The procedure was carried out according to the Invitrogen Lipofectamine 3000 Transfection Reagent Kit. Simply put, cell culturing was done at 37℃ with a CO_2_ incubation confluency of 70 to 80%. In two 1.5 mL EP tubes, the following two solutions were prepared separately: Tube A: 125 µL opti-MEM + 5 µL Lipo3000. Tube B: 125 µL opti-MEM + 1-2.5 µg DNA (1 ug pCKK002 + 1 µg pCKK005 + 1 µg pCKK006 + 2 µg FDX1 donor DNA fragment) + 5 µL P3000. The liquid from tube A was added to tube B, and it was gently mixed and incubated at room temperature for 10 to 15 min. The DNA-lipid complex was then added to the cells. The plate was gently shaken to ensure the even distribution of the complex across all wells, and the cells were incubated at 37℃ for 2 days. Then, cells were taken, digested with 0.25% trypsin, and blown into individual cells, and the cells were suspended in 10% fetal bovine serum DMEM medium (400 µg/mL G418, with a final insulin concentration of 10 μg/ml,) for later use. The cell suspension was diluted in gradient multiples and seeded into a 96-well culture plate containing 10 mL of 37℃ pre-warmed DMEM medium (containing 10% FBS) at gradient densities of 5, 10, 20, and 50 cells per well, and it was gently rotated to disperse the cells evenly. It was then placed in a cell incubator with 37℃, 5% CO_2_, and saturated humidity for 2 to 3 weeks.

### Cell Roliferation/Toxicity Assay

The cells were seeded at a density of 1 × 10^4^ cells per well in 96-well plates. After 24 hours of incubation to allow for cell adhesion and growth, the cells were treated with 100 μL of DMEM medium containing serial concentrations of Fu-Cu (0, 40, 80, 160, 200, 320, and 400 μM), CuCl_2_ (0, 40, 80, 160, 200, 320, and 400 μM), and Fu (0, 15, 30, 60, 75, 120 and 150 μg/mL) for an additional 24 hours. The Cell Proliferation/Toxicity Assay was conducted using the CCK-8 method (Abbkine, Wuhan, China). Initially, cells were seeded at a density of 5,000 cells per well in a 96-well plate and incubated for 24 hours at 37℃ in a 5% CO_2_ environment. Following this, the test substance was introduced at various concentrations and the plate was incubated for an additional 24 hours. Subsequently, 10 μL of CCK-8 solution was carefully added to each well, ensuring no bubbles were introduced that could interfere with the optical density (OD) reading. After a further 1.5-hour incubation, adjusted based on cell type and density, the absorbance at 450 nm was measured using a microplate reader.

### HuH-7 Transwell Invasion and Migration Assays

For transwell invasion and migration assays, cells were grown to a density of 70%-80%, followed by incubation in DMEM (containing 10% FBS) medium within 6-well plates (Corning Costar Corp) as per the manufacturer’s guidelines. After a 24 h incubation period, cells were harvested using EDTA-free trypsin (Gibco), collected, and resuspended in DMEM medium, with the addition of Null, Fu-Cu (320 μM), Fu (120 μg/mL), CuCl₂ (320 μM), and Es-Cu (100 nM elesclomol + 10 μM CuCl₂), adjusting the cell density to 2.5 × 10^5^. For migration assays, 200 μL of the cell suspension was placed into the upper chambers of a 24-well transwell plate containing a PC membrane (Corning Costar) and incubated for 24 h. Conversely, for invasion assays, a 1:8 dilution of Gold Medal Matrix Gel (ABW) was added to the upper chambers and incubated at 37°C for 3 hours to solidify before seeding the cells. The lower chambers were filled with 500 μL of medium supplemented with 15% FBS. Following a 24 hours incubation period at 37°C, cells that migrated or invaded through the membrane were imaged and counted in three distinct fields under a 200 × objective lens using a microscope.

### Cell copper content Assays

HuH-7 cells in the logarithmic growth phase were divided into the following experimental groups: Null, Fu-Cu (320 μM), Fu (120 μg/mL), CuCl₂ (320 μM), and Es-Cu (100 nM elesclomol + 10 μM CuCl₂). After 24 hours of treatment, the cells were collected, and Cu²⁺ ion content was measured using a commercially available Cell Copper Content Assay Kit (Solarbio, BC5755), following the manufacturer’s protocol. Total copper content in HuH-7 cells was also determined using ICP-MS. Cells were treated with Fu-Cu (320 μM) for various durations (0, 4, 8, 12, and 16 hours), and copper levels were measured at each time point.

### TEM imaging of Fu-Cu in HuH-7 cells

Approximately 10^7^ cells treated with 320 μM for 24 hours were immediately fixed in 1.5 mL of 2.5% glutaraldehyde in 2 mL centrifuge tubes and placed in a 4°C refrigerator for 12-24 hours. Samples were then post-fixed in 1% osmium tetroxide solution for 1-2 hours. The osmium tetroxide was carefully removed, and samples were washed three times in 0.1M phosphate buffer (pH 7.4) for 15 minutes each. Samples underwent a gradient dehydration in 30%, 50%, 70%, 80%, and 95% acetone for 10 minutes each, followed by two rounds of dehydration in 100% acetone (2 × 20 min). For infiltration and embedding, the samples were subjected to a mixture of acetone and embedding medium at ratios of 3:1 at 37°C for 1 hour and 1:1 at 37°C for 3 hours, followed by pure embedding medium at 37°C overnight. The samples were then embedded in resin blocks and placed in a 60°C oven for polymerization for 48 hours. Then, 70-90 nm thick, were cut using an ultramicrotome and collected on copper grids. Staining was carried out with uranyl acetate for 8-15 minutes and lead citrate for 5-10 minutes, before drying. The samples were finally examined and images were captured using TEM.

### Apoptosis Analysis in Cells

HuH-7 cells in the logarithmic growth phase were collected and adjusted to a cell density of 5 × 10⁵ cells/mL. A volume of 2 mL per well was evenly seeded into 6-well plates and incubated at 37°C with 5% CO₂ until cell confluence reached approximately 80%. The culture medium was removed, and cells were gently rinsed with PBS. Experimental groups were treated with the Null, Fu-Cu (320 μM), Fu (120 μg/mL), CuCl₂ (320 μM), and Es-Cu (100 nM elesclomol + 10 μM CuCl₂) for 12 hours and 24 hours. Additionally, the drug treatment times for the Null group and the Fu-Cu group were extended by 6 hours and 18 hours, respectively. Apoptosis was assessed using the Annexin V-FITC/PI Apoptosis Detection Kit (40302ES60, Yeason) according to the manufacturer’s instructions. Briefly, cells were harvested using 0.25% trypsin without EDTA and collected by centrifugation. The cells were washed twice with pre-cooled PBS and resuspended in 100 µL of 1× Binding Buffer by gentle pipetting. Under light-protected conditions, 5 µL of Annexin V-FITC and 10 µL of PI staining solution were added and mixed gently. After incubation in the dark for 10 minutes, 400 µL of 1× Binding Buffer was added, mixed gently, and the samples were immediately analyzed. Single-stained controls with Annexin V-FITC or PI were prepared to adjust for compensation.

### ROS Detection of cells

Cells were treated with Null, Fu-Cu (320 μM), Fu (120 μg/mL), CuCl₂ (320 μM), and Es-Cu (100 nM elesclomol + 10 μM CuCl₂) for 18 hours, following the protocol outlined in the Reactive Oxygen Species Assay Kit (Biosharp, BL714A). H_2_DCFDA was diluted to a final concentration of 10 µM in DMEM medium at a ratio of 1:1000 for ROS probe preparation. For probe loading, cells were harvested and resuspended in the diluted probe solution at a cell density of 1 × 10⁶ to 2 × 10⁷ cells/mL, followed by incubation in the dark for 30 minutes at 37°C in a cell incubator. After incubation, cells were washed 1 to 2 times with DMEM medium to remove unincorporated H_2_DCFDA. Flow cytometry analysis was performed using the FITC channel, with excitation at 488 nm and emission at 530 nm. Data were analyzed using FlowJo software, and the mean fluorescence intensity (MFI) was calculated and plotted.

### GSH-Responsiveness of Fu-Cu nanoparticles

The HuH-7, LX-2, and HEK293A cell lines were seeded in 6-well culture plates and allowed an initial 24 hours incubation period to permit adherence and proliferation. The cells were then treated with Null, Fu-Cu (320 μM), Fu (120 μg/mL), CuCl₂ (320 μM), and Es-Cu (100 nM elesclomol + 10 μM CuCl₂) dissolved in 1.5 mL DMEM per well for 4 hours. Reduced glutathione (GSH) test kit (Solarbio, BC1175) was used to assay GSH level. Briefly, we collected 1 × 10^7^ cells and split the supernatant. The measurements were carried out according to the manufacturer’s protocols. The microplate reader was calibrated to detect at a wavelength of 412 nm.

### Copper-ATPase assay

Cells treated with the Null, Fu-Cu (320 μM), Fu (120 μg/mL), CuCl₂ (320 μM), and Es-Cu (100 nM elesclomol + 10 μM CuCl₂) were collected and lysed by adding 100 µL of cell lysis buffer (containing PMSF) per 1 × 10⁶ cells. After thorough lysis, the lysate was centrifuged at 10,000 rpm for 10 minutes at 4°C, and the supernatant was collected for further analysis. Reagent preparation and sample measurements were performed according to the instructions of the Copper-ATPase assay kit (A057-1-1, Jiancheng).

### Bioinformatics Analysis of HCC

Transcriptomic data for 371 HCC patients, alongside 276 healthy individuals, were sourced from the TCGA (The Cancer Genome Atlas, https://portal.gdc.cancer.gov) repository, presented in HTseq-FPKM format. To facilitate downstream analyses, the raw expression values were subjected to a log2 transformation, converting them to TPM format. A set of genes implicated in cuproptosis was delineated based on insights from two recent publications^5,31^. These gene set includes *SLC31A1*, *ATP7A*, *ATP7B*, *ATOX1*, *LOX*, *SCO1*, and others. Detailed data can be found in Supplementary Tables S5 and S6. The log-rank test was used to compare survival differences between the two groups in the Kaplan-Meier (KM) survival analysis. The p-value, hazard ratio (HR), and 95% confidence interval (CI) were calculated using the log-rank test and univariate Cox regression. Detailed data are provided in Supplementary Tables S7. Analytical procedures and statistical methodologies were executed using R software, version 4.0.3.

### Caenorhabditis elegans Culture and RNAi

Wild-type N2 Bristol *C. elegans* were sourced from the Caenorhabditis Genetics Center. These worms were cultivated at 20℃ on nematode growth medium (NGM) agar plates and nourished with an *E. coli* OP50 lawn. Strains used in our study are listed in Supplementary Table S4.

The gene CDS fragments of the target genes was integrated into the L4440 vector^40^, which was pre-configured with the T7 Promoter and 2 enzyme restriction sites (*Hind*III and *BamH*I), utilizing the Gibson cloning technique. Subsequently, either RNAi plasmids or an empty vector were transformed into *E. coli* HT115 (DE3) bacterial strains. A selected colony was inoculated in 6 mL of LB medium supplemented with Ampicillin (100 μg/mL) and incubated at 37°C overnight. The cultured bacteria were then harvested in 600 μL of LB, and a fraction (1/6) was spread onto the center of an LB agar plate, also containing Ampicillin (100 μg/mL). The plates were allowed to dry and were wrapped in aluminum foil for overnight induction at room temperature. L4 larval worms were isolated and nourished with *E. coli* HT115 (DE3) bacteria transformed with the dsRNAi-containing plasmid DNA. After feeding, worms were selected for further analysis, including genotyping or RT-qPCR confirmation.

### Microinjection and Transgenic Lines of Overexpression Worms

To generate plasmid encoding *ccs*-208 and *mtch2*-201 used for tissue expression, cloned into pPD95.75 at the *Xma*I and *EcoR*I sites^41^. To generate transgenic lines (Pdpy-30:: *ccs*-208 and Pdpy-30::*mtch2*-201), the constructs (*ccs*-208 and *mtch2*-201 in pPD95.75) were modified by placing dpy-30 promoter upstream of the *ccs*-208 and *mtch2*-201 coding sequence. The sequences of human *MTCH2* and *CCS*-specific nested PCR primers used for the 3’ RACE analysis are listed in Supplementary Table S3.

Gonad microinjection was performed according to standard *C. elegans* procedures in young adult hermaphrodites. To create transgenic lines, 20 ng/μL of pCKK041 (P*dpy-30*::*ccs-208*) and pCKK042 (P*dpy-30*::*mtch2-201)* plasmids, along with 20 ng/μL of pCKK043 (P*col-10*::*sfGFP*) plasmid as a co-injection transgene marker, were injected into each gonad. At least three independent lines, stably carrying plasmid DNA as extra chromosomal arrays, were obtained per construct. The transgenic lines used in our study are listed in Supplementary Table S8.

### *C. elegans* Toxicity Assay

S medium was supplemented with Fu-Cu at concentrations of 0, 1200 and 2400 μM. Plates were seeded with *E. coli* (either OP50 or HT115 (DE3)) three days before the addition of synchronized *C. elegans* L1 larvae. The details of S medium can be found in Supplementary Table S9. 90-100 adult worms were examined for viability 48 hours post-plating. A two-tailed student’s *t*-test was used to compare sterility rates between control and experimental populations.

For dsRNA-treated worms, wild-type hermaphrodites were transferred to RNAi plates during young, gravid adulthood^42^. *E. coli* HT115 (DE3) used for RNAi feeding were grown to an OD_600_ of 0.5 in LB medium supplemented with 50 μg/mL ampicillin. L4/young adult stages were transferred to plates with 100 µg/mL FUdR, 50 μg/mL ampicillin, and 0.4 mM IPTG. Empty vectors (L4440) served as controls.

### C. elegans ROS Assay

Synchronized L4 stage nematodes were transferred to NGM plates containing 0 and 2400 μM Fu-Cu (with 100 μg/mL FUdR) and incubated at 20°C for 5 days. The nematodes from each group were washed and collected using M9 buffer, transferred to 1.5 mL centrifuge tubes, and centrifuged at 3, 000 rpm for 2 minutes. This washing step was repeated at least 3 times until the nematodes were clear and individually visible. After washing, the supernatant was discarded, and 100 μL of M9 buffer was retained. Then, 1.5 μL of 10 mM DCFH-DA working solution was added to achieve a final concentration of 150 μmol/L (Beyotime, ROS Assay Kit, S0033S). The samples were incubated at 37°C with shaking at 100 rpm, in the dark, for 2 hours. After staining, 1 mL of M9 buffer was added and centrifuged at 3, 000 rpm for 2 minutes. The supernatant was discarded, and this washing step was repeated at least 3 times to thoroughly clean the probe. An appropriate number of nematodes was then transferred to a 2% agarose pad, covered with a coverslip, and images were captured under the FITC green fluorescence channel.

### Establishment of Tumor Models

Female athymic BALB/c nude mice, aged 6-8 weeks, were procured from Vital River Laboratories in Beijing, China, and maintained under specific-pathogen-free conditions at the Laboratory Animal Center of Jennio Biotech Co., Ltd. All animal procedures were conducted in compliance with the guidelines set forth by the Institutional Animal Care and Use Committee of Jennio Biotech Co., Ltd. A HuH-7 tumor-bearing mouse model was established by subcutaneously injecting 1 × 10^7^ HuH-7 cells in a 150 μL suspension, combined at a 1:1 ratio with Matrigel, into the left flank of the mice.

### In *vivo* Antitumor Efficacy Study

Once the tumors reached an approximate volume of 100 mm^3^, the mice were randomly allocated into five groups, each containing six mice: PBS, CuCl_2_ (3 mg/kg), Fu (165 mg/kg), Cu-Es (1.5 mg/kg CuCl_2_ +6.25 mmol/kg elesclomol) and Fu-Cu (87.5 mg/kg). Intravenous injections of either the designated nanomaterial solutions at a concentration of 10 mg/kg or PBS were administered on days 0 and 7. Tumor volume and body weight were monitored every three days, with tumor volume (V) calculated using the formula V = (Width^2^ × Length)/2. After three weeks, the mice were euthanized, and photographs were taken. The tumors were excised, photographed, and their final weights recorded.

### H&E and TUNEL Staining

Upon concluding the treatment, tumor tissues from the mice were isolated and fixed in a 10% paraformaldehyde solution. Following standard tissue preparation, H&E assay was conducted and the samples were observed under a Nikon Eclipse E100 optical microscope. Apoptotic cells were detected using a TUNEL assay, facilitated by an *in situ* apoptosis detection kit (Servicebio, China), in accordance with the manufacturer’s guidelines. Nuclei were stained with DAPI (Servicebio, China), and an anti-fluorescence quenching solution (Servicebio, China) was subsequently applied. Fluorescent images were captured using a Nikon Eclipse C1 microscope.

### Western Blotting Analysis

Following separation on a 10% SDS–PAGE gel, protein extracts were transferred to a polyvinylidene fluoride (PVDF) membrane. The membrane was then blocked with 5% skim milk at room temperature for 1 hour. Primary antibodies were added to the membrane, which was incubated overnight at 4°C. Subsequently, the membrane was incubated with secondary antibodies. The specific antibodies utilized are detailed in Supplementary Table S10. Protein bands were visualized using a Tanon-5200 Chemiluminescent Imaging System (Tanon, China). The complete, uncut original images are provided in Figure S7.

### Immunofluorescence Analysis

The cells were fixed using 3.7% formaldehyde and permeabilized with 0.1% Triton X-100. Following two washes with PBST (1× PBS containing 0.1% Tween-20), 50 μl of blocking solution (1× PBS, 1% BSA, 0.1% Tween-20) was applied. Subsequently, the cells were incubated with the primary antibody overnight at 4°C, followed by incubation with the conjugated secondary antibody and DAPI at room temperature. Imaging was performed using the LSM710 confocal microscope (Zeiss, Pleasanton, CA, USA).

Fresh tissue was fixed in 4% paraformaldehyde for over 24 hours, trimmed, and placed in a dehydration cassette. The tissue was dehydrated through a graded ethanol series (65%, 75%, 90%, 95%, and 100%) and cleared with a clearing agent, followed by paraffin infiltration and embedding. Sections were cut at 3-4 μm, floated on 42°C water, mounted on slides, and dried at 65°C. After deparaffinization and rehydration, antigen retrieval was performed in citrate buffer under high pressure. Endogenous peroxidase activity was blocked with 3% H_2_O_2_, and sections were blocked with 5% BSA. Primary antibodies were incubated overnight at 4°C or for 1 hour at room temperature, followed by secondary antibody incubation for 1 hour. Nuclei were counterstained with DAPI, and slides were mounted with antifade medium. Finally, slides were scanned using a 3D Pannoramic MIDI scanner. The details of all antibodies utilized are provided in Supplementary Table S10.

### Flow Cytometry Analysis

Peripheral blood mononuclear cells were isolated and resuspended in 1 × PBS after thawing. Cells were divided into two equal portions and transferred to flow cytometry tubes, with a small aliquot retained as the blank control. Cells were centrifuged at 1500 rpm for 5 min at room temperature, and the supernatant was discarded. For staining, 30 μL of diluted antibody mix was added to the cell pellet and gently mixed. The cells were incubated for 30 min at room temperature or 4°C. Following staining, 500 μL of 1 × PBS was added directly to each tube, and the cells were centrifuged again at 1500 rpm for 5 min at room temperature. The supernatant was carefully aspirated, and the cells were resuspended in 200 μL of 1 × PBS for analysis on an ACEA NovoCyte flow cytometer. The following antibodies and detection channels were used: CD3 (AF700), CD19 (PE-CY7), and Ly6G (APC-CY7).

### RNA extraction and RT-qPCR assays

Total RNA was extracted from HuH-7 cells and *C. elegans* utilizing TRIzol Reagent (Takara Biotechnology, China). The extracted RNA was then reverse transcribed to cDNA using the PrimerScript RT-PCR kit (Takara Biotechnology, China). Quantitative real-time PCR (RT-qPCR) was performed employing a SYBR Green reaction mix (Qiagen, Germany) on a LightCycler 96 System (Roche). The expression levels of *GAPDH* or *β*-*actin* genes were used as internal controls for normalization. All primers utilized in this study are detailed in Supplementary Table S3.

### Statistical Analysis

Data are represented as mean ± standard deviation (SD) for a sample size (n) ranging from 3 to 5, unless specified otherwise. For the analysis of cuproptosis relative assays, the viability of co-treated cells was denoted as a fold change compared to cells without drugs treatment. Statistical evaluations were conducted using GraphPad Prism version 9 (GraphPad Software, San Diego, CA). The significance of differences between multiple groups was determined through ONE-way ANOVA, followed by Bonferroni’s post-hoc test.

## Acknowledgements

The HEK293A cell line was gifted from the laboratory of Dr. Guoqin Liu in SCUT. This project was supported by the National Natural Science Foundation of China (32372286 and 32072201), Guangdong Basic and Applied Basic Research Foundation (2023A1515011967 and 2023A1515012223).

## Supplementary information

In separated documents including supplementary information and tables.

## Data availability

Data are available from the corresponding author upon reasonable request.

## Author contributions

H.C. was the main author of this work. H.C. & X.P., conducted the wet-lab experiments, data analysis, visualization, wrote the original draft; Q.W. conducted the *C. elegans* experiments, data analysis, visualization; Z.C. performed the bioinformatics analysis; L.Z., J.L., Q.Z., R.Z. & X.Z. conducted some experiments; L.Y. & L.Z. revised the manuscript. J.J. provided an overall assessment and acquired funding of this work. All authors reviewed and provided input on the manuscript.

