## Supplementary Material for "Fucoidan-Copper Nanoparticles to Potentiate Synergistic Cancer Cell Cuproptosis and Immunotherapy"

#### Supplementary Tables

**Table S1** Copper element content of Fu-Cu nanoparticles.

**Table S2** Reagents and chemicals used in this study.

**Table S3** All Oligos/primers used in this study.

**Table S4** All plasmids used in this study.

**Table S5** Cuproptosis-Related Transcriptomic Patterns in 371 HCC Patients Versus 276 Healthy Individuals.

**Table S6** Survival statistics of HCC patients with high and low *MTCH2* expression.

**Table S7** Statistical data of *MTCH2* expression at different stages in HCC patients.

**Table S8** Genes utilized for the construction of *C. elegans* strains in this study.

**Table S9** S medium containing for cultivation of *C. elegans.*

**Table S10** List of antibodies used for immunostaining, western blotting and flow cytometry.

**Supplementary note 1, Hematological Evaluations for Toxicology Evaluation**

In the study, BALB/c mice (n = 3 per treatment group) received intravenous injections of either PBS, $CuCl_2$, fucoidan, Es-Cu or Fu-Cu. Blood glucose levels were continuously monitored for a 48-hour period post-injection using a commercial glucometer. At the 20-hour mark post-injection, the mice were euthanized for blood collection to perform hemogram and blood biochemistry analyses. Hemoglobin (HGB), white blood cells (WBC), and platelets (PLT) were quantified using a Mindray BC-2800Vet Automated Haematology Analyzer (China). Plasma levels of alanine aminotransferase (ALT), aspartate aminotransferase (AST), lactate dehydrogenase (LDH), and creatinine (CR) were assessed using a Chemray 800 Automatic Biochemical Analyzer (China). Levels of TNF-$\alpha$ and IL-6 were determined using ELISA kits, following the manufacturer's guidelines.

### Supplementary Figures

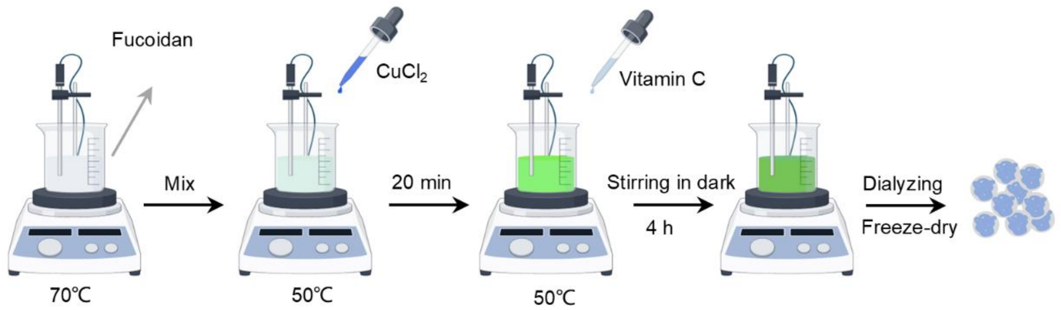


**Figure S1. Schematic representation of the synthesis of Fu-Cu nanoparticles**. Fucoidan solution was mixed and heated to 70°C, followed by the addition of $CuCl_2$ at 50°C with continuous stirring for 20 minutes. Subsequently, Vitamin C was added to the mixture, and the reaction was stirred in the dark for 4 hours at 50°C. The resulting solution was dialyzed and then freeze-dried to obtain the final Fu-Cu nanoparticle product.

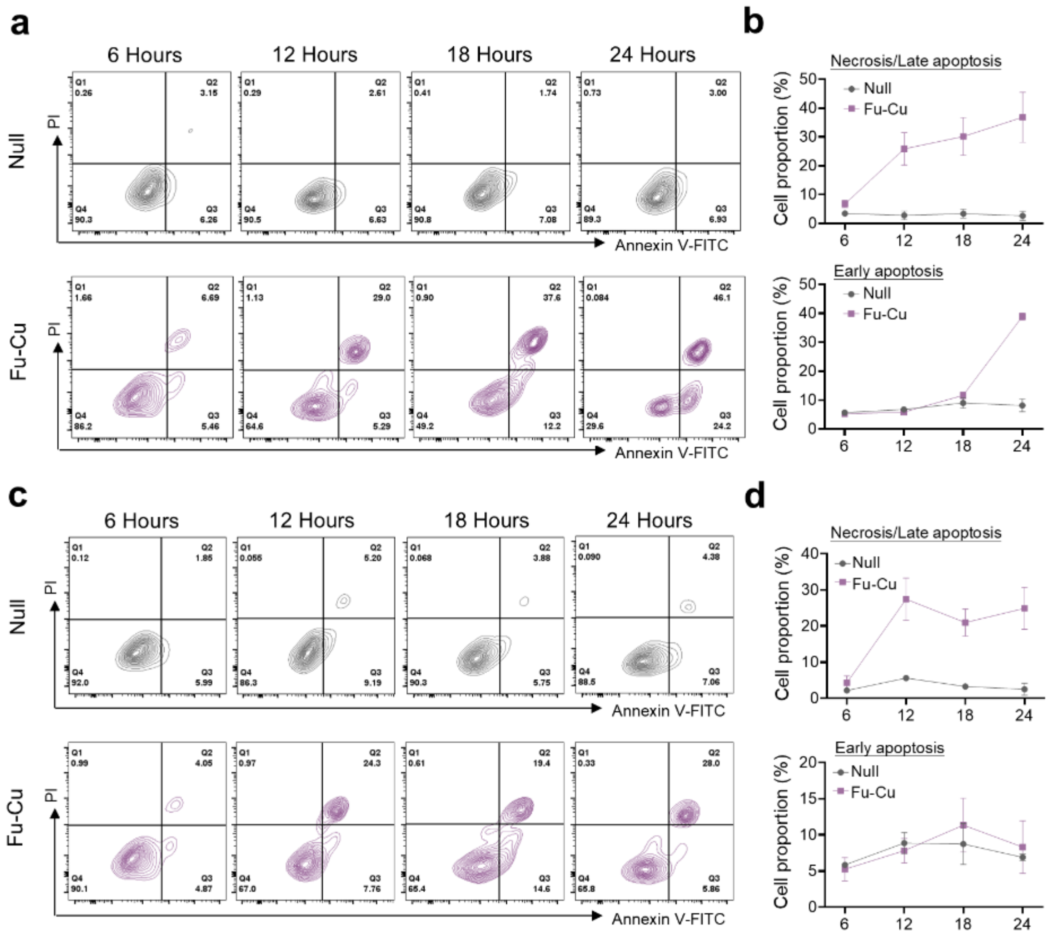


**Figure S2. Fu-Cu induces apoptosis in HuH-7 cells over time.** (**a**) Flow cytometry analysis of Annexin V-FITC and PI staining in HuH-7 cells treated with either the Null or Fu-Cu (320 μM) for 6, 12, 18, and 24 hours. (**b**) Quantification of necrosis/late apoptosis and early apoptosis in HuH-7 cells treated with Fu-Cu compared to Null at the indicated time points. (**c**) Flow cytometry analysis of Annexin V-FITC and PI staining in HuH-7 *FDX1* KO cells treated with either the Null or Fu-Cu (320 μM) for 6, 12, 18, and 24 hours. (**d**) Quantification of necrosis/late apoptosis and early apoptosis in LX-2 cells treated with Fu-Cu compared to Null at the indicated time points. The plots show the distribution of early apoptotic cells (Annexin V$^+$/PI$^-$), late apoptotic/necrotic cells (Annexin V$^+$/PI$^+$), and viable cells (Annexin V$^-$/PI$^-$). Data represent mean ± SD (n = 3).

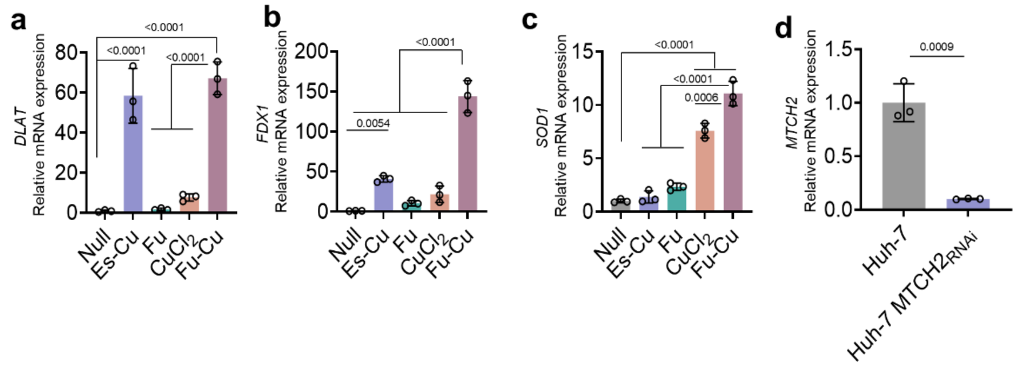


**Figure S3. Fu-Cu treatment induces cuproptosis-related gene expression in HuH-7 cells.** Relative mRNA expression levels of *DLAT* (**a**), *FDX1* (**b**), and *SOD1* (**c**) in HuH-7 cells after treatment with Null, Es-Cu (100 nM elesclomol + 10 µM $CuCl_2$), Fu (120 µg/mL), $CuCl_2$ (320 µM), or Fu-Cu (320 µM). (**d**) Relative mRNA expression of *MTCH2* in HuH-7 cells compared to HuH-7 cells treated with *MTCH2* siRNA. Gene expression was normalized to *β-actin*, and data are shown as mean ± SD (n = 3). Statistical significance was determined by one-way ANOVA, with p-values indicated in the graphs.

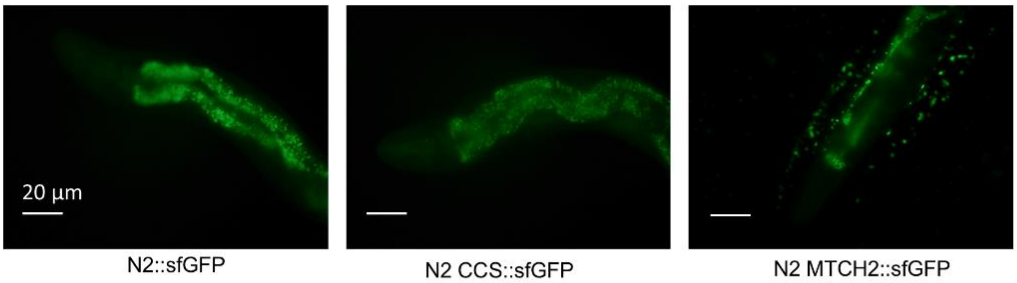


**Figure S4. Fluorescence imaging of different *C. elegans* strains expressing sfGFP-tagged proteins.** Representative fluorescence microscopy images of *C. elegans* strains N2::sfGFP (control), N2 CCS::sfGFP, and N2 MTCH2::sfGFP, showing the localization of sfGFP, CCS::sfGFP, and MTCH2::sfGFP, respectively. Scale bar = 20 µm.

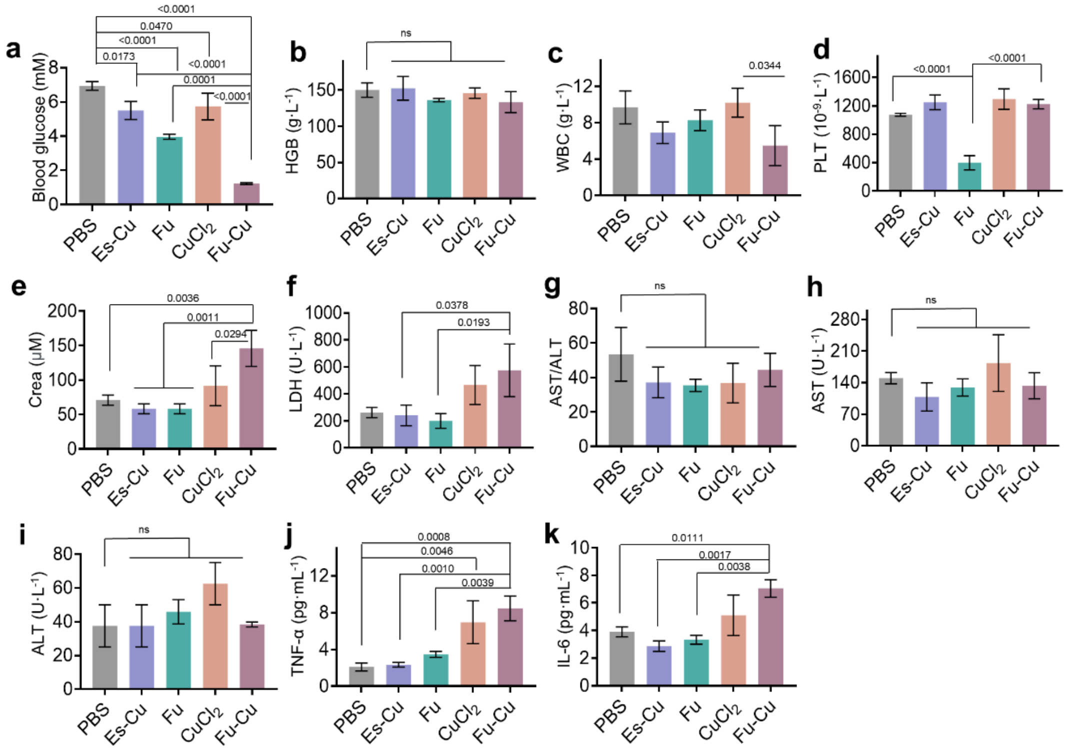


**Figure S5. Effect of Fu-Cu and other treatments on blood glucose, hematological parameters, and biochemical markers in BALB/c mice.** (**a**) Blood glucose levels measured 20 hours after intravenous injection of PBS, Es-Cu, Fu, $CuCl_2$, or Fu-Cu. Blood glucose significantly decreased in the Fu-Cu group compared to other treatments. Hemogram analysis showing levels of hemoglobin (HGB) (**b**), white blood cells (WBC) (**c**), platelets (PLT) (**d**), and creatinine (Crea) (**e**) after 20 hours. Blood biochemistry analysis showing lactate dehydrogenase (LDH) (**f**), AST/ALT ratio (**g**), aspartate aminotransferase (AST) (**h**), and alanine aminotransferase (ALT) (**i**) levels across the treatment groups. The Fu-Cu group showed elevated LDH levels. Plasma levels of pro-inflammatory cytokines TNF-α (**j**) and IL-6 (**k**) measured by ELISA. Fu-Cu treatment significantly increased TNF-α and IL-6 levels compared to other groups. Data represent mean ± SD (n = 3 per group), with statistical significance determined by one-way ANOVA.

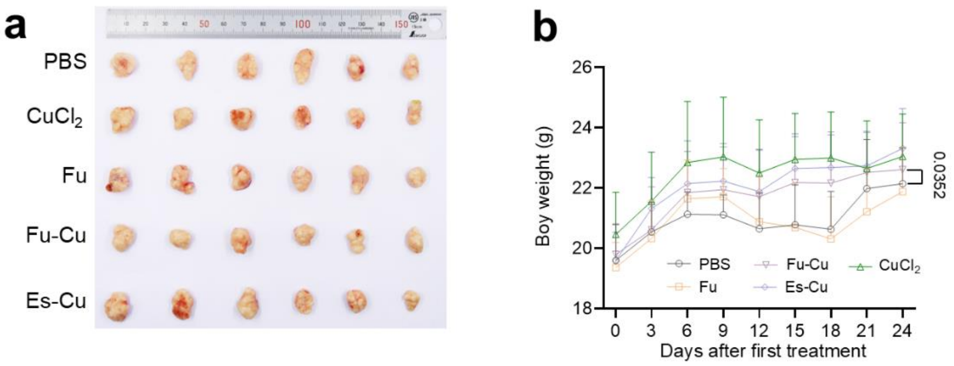


**Figure S6. Fu-Cu treatment inhibits tumor growth in a BALB/c nude mice subcutaneous tumor model.** (**a**) Images of excised tumors from BALB/c nude mice treated with PBS, $CuCl_2$, Fu, Fu-Cu, or Es-Cu. Tumors were collected at the end of the experiment to assess the effect of different treatments on tumor growth. (**b**) Body weight of BALB/c nude mice was monitored over 24 days after the first treatment. Mice received intravenous injections of PBS, $CuCl_2$, Fu, Fu-Cu, or Es-Cu. Data represent mean ± SD (n = 6 per group).

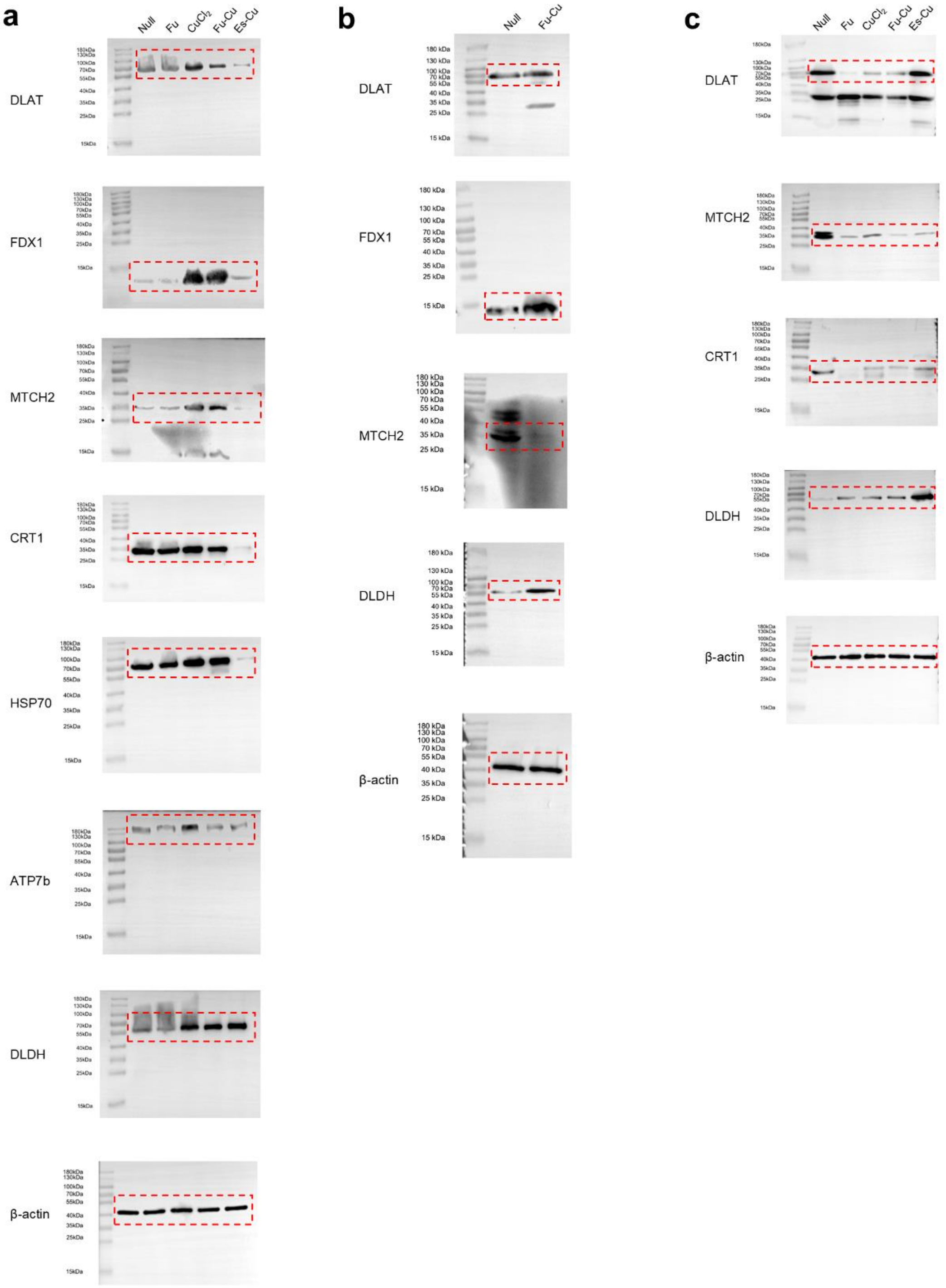


**Figure S7. Uncropped Western blots for Figure 3f (a), Figure 3g (b), and Figure 4i(c).**
